# Clinical PARP inhibitors allosterically induce PARP2 retention on DNA

**DOI:** 10.1101/2022.11.08.515634

**Authors:** Marie-France Langelier, Xiaohui Lin, Shan Zha, John M. Pascal

**Affiliations:** Department of Biochemistry and Molecular Medicine, Université de Montréal, Montréal, QC H3C 3J7 Canada; Institute for Cancer Genetics, Vagelos College for Physicians and Surgeons, Columbia University, New York City, NY 10032 USA

## Abstract

PARP1 and PARP2 detect DNA breaks, which activates their catalytic production of poly(ADP-ribose) that recruits repair factors and releases PARP1/2 from DNA. PARP inhibitors (PARPi) are used in cancer treatment and target PARP1/2 catalytic activity, interfering with repair and increasing PARP1/2 persistence on DNA damage. Additionally, certain PARPi exert allosteric effects that increase PARP1 retention on DNA. However, no clinical PARPi exhibit this allosteric behavior toward PARP1. In contrast, we show that certain clinical PARPi exhibit an allosteric effect that retains PARP2 on DNA breaks in a manner that depends on communication between the catalytic and DNA-binding regions. Using a PARP2 mutant that mimics an allosteric inhibitor effect, we observed increased PARP2 retention at cellular damage sites. The new PARPi AZD5305 also exhibited a clear reverse allosteric effect on PARP2. Our results can help explain the toxicity of clinical PARPi and suggest ways to improve PARPi moving forward.

## Introduction

PARP1 and PARP2 are members of the PARP family of proteins that includes 17 members involved in a variety of cellular processes including gene transcription, chromatin regulation, the antiviral response, and cellular signalling (*1*). PARP1 and PARP2 use NAD^+^ as a substrate to create poly(ADP-ribose) (PAR) covalently attached to target proteins and DNA (*2*). PARP1 and PARP2 play critical roles in DNA repair by detecting DNA damage and signaling the presence of a DNA break through the production of PAR, leading to recruitment of repair factors to the site of damage. Recently, histone PARylation factor 1 (HPF1) has been identified as an important regulator of PARP1 and PARP2 activity in the response to DNA damage (*3*–*5*). HPF1 switches the target residues modified by PARP1 and PARP2 from aspartate/glutamate to serine by inserting a catalytic glutamate in the active site of PARP1 and PARP2 to allow serine deprotonation and subsequent ADP-ribosylation (*6, 7*). HPF1 also changes the preference of substrate from *cis* modification to *trans* modification and stimulates the initiation steps of the ADP-ribosylation process, while blocking or limiting the growth of PAR by occupying the ADP-ribose elongation site (*3, 8, 9*).

PARP1 is a modular protein composed of seven distinct domains (Fig. 1). The Zn1 and Zn2 domains recognize and bind to the DNA break (*10, 11*). The Zn3 and WGR domains also contribute to DNA binding (*12*). An automodification domain is composed of a linker region that holds the main residues targeted for modification (*13, 14*), and a BRCT fold that was recently reported to contribute to PARP1 interaction with undamaged DNA (*15*). The CAT domain consists of two sub-domains, the helical domain (HD) and ADP-ribosyltransferase (ART) domain, which is conserved in all PARP family members (*1*). PARP2 is a smaller protein, with a short unstructured N-terminal region, a WGR, and a CAT domain. PARP2 relies mostly on the WGR domain for DNA binding (*16*–*19*). We have shown that the HD of PARP1 blocks the access of NAD^+^ to the active site (*20, 21*). In recognizing DNA breaks, the regulatory domains of PARP1 (Zn1, Zn3 and WGR) assemble on the break and form a binding platform for the HD (*12*). This multi-domain assembly leads to a local destabilization of the HD that opens the active site so that NAD^+^ can bind (*12, 20, 21*). This substrate blocking mechanism is conserved in PARP2 and PARP3 (*21*). Recently, we have captured the structure of a PARP1 HD mutant that favors the active conformation that is open for NAD^+^ binding and forms an extended interface with WGR (*22*). In this active conformation, the HD contributes to PARP1 multi-domain assembly and retention on DNA (*22*).

**Fig. 1.**
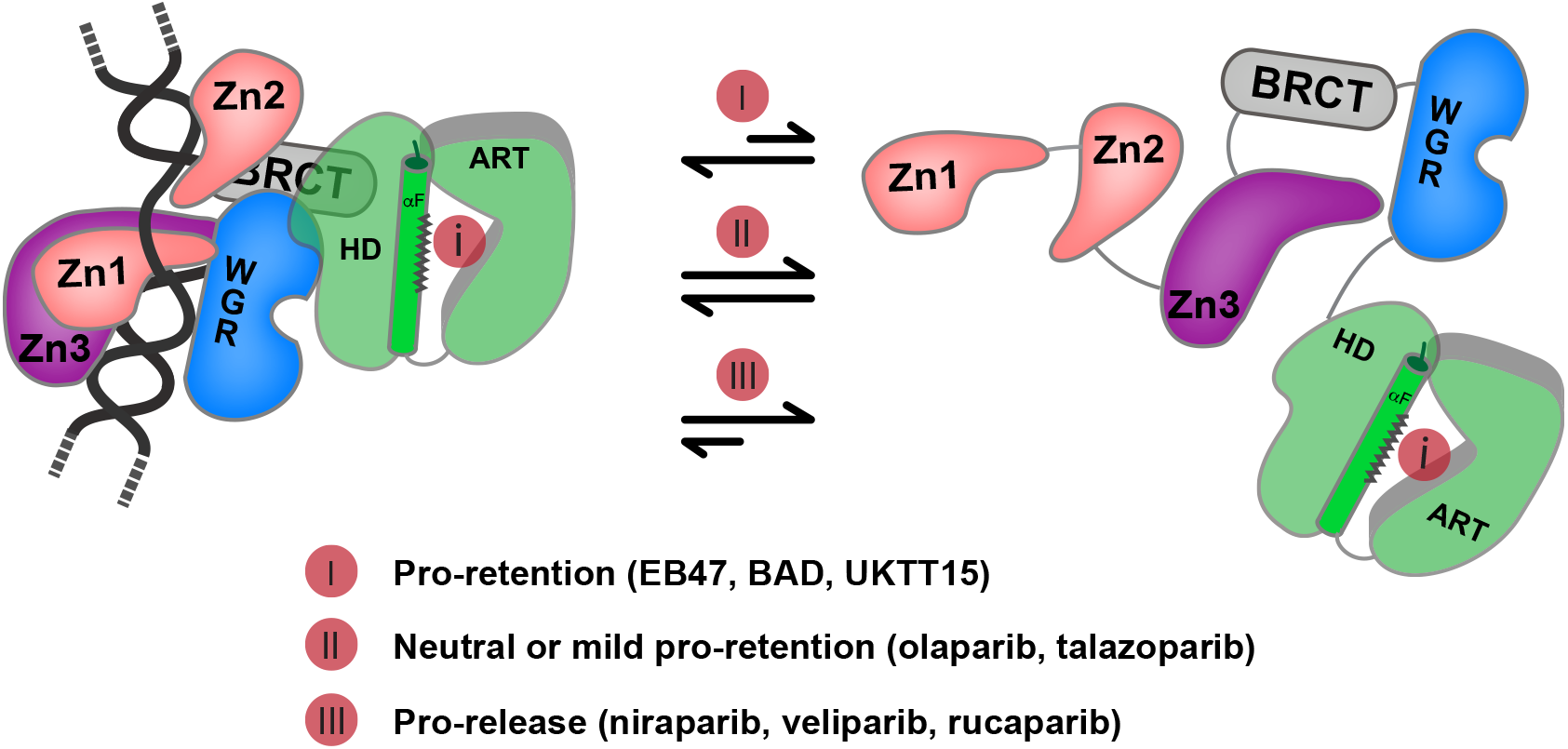
PARPi classification based on reverse allostery in PARP1. PARPi classify in 3 different types based on their ability to modulate PARP1 retention on DNA through allosteric effects. Type I inhibitors have a pro-retention effect, type II have no effect or mild pro-retention effect, and type III have a pro-release effect (*33*). The length of the arrows indicate the distribution towards DNA retained PARP1 on the left and released PARP1 on the right.

PARP inhibitors (PARPi) have been developed extensively since the discovery that they can selectively kill BRCA deficient cancer cells, which are defective in DNA repair, a phenomenon called synthetic lethality (*23, 24*). They are broadly used in the clinic to treat various forms of cancer (*25*). However, recently there has been a push to develop novel and more selective PARP1 inhibitors that do not target other PARP family members. Indeed, the ability of PARPi to kill cancer cells has been attributed to PARP1 trapping on DNA (*26, 27*) and the non-specific targeting of PARP2 was suggested to be responsible for some of the hematological side effects observed with the current clinical PARPi (*28*). PARP1 trapped on DNA acts as a toxic lesion that blocks the progression of the replication fork leading to the creation of double-strand breaks that are particularly lethal in BRCA deficient cells, which are impaired in their ability to perform homologous recombination (*29*). The ability of PARPi to trap PARP1 on DNA varies between inhibitors and was proposed to be related to the propensity of some PARPi to increase PARP1 retention on DNA independently of catalytic inhibition (*26, 30*). This mechanism was termed reverse allostery since PARPi bind the catalytic site and increase the ability of the regulatory domains to bind DNA. Other studies have suggested that the differences in the ability of PARPi to trap PARP1 and kill cancer cells are only related to inhibitory potencies and variations in inhibitor off rates (*31, 32*).

Recently, we have analyzed the ability of PARPi to induce reverse allostery in PARP1 and classified them in 3 different types (*33*) (Fig. 1). Type I PARPi have a pro-retention allosteric effect. They exert this pro-retention effect by increasing the dynamics of HD helices and favoring the HD conformation that interacts with WGR and increases DNA affinity (*33*). The NAD^+^ analog BAD, EB47, and the veliparib derivative UKTT15 are type I PARPi, but none of the clinical inhibitors belong to this class in PARP1 (*21, 33*). Type II PARPi have no effect or a mild effect on PARP1 allostery and retention on DNA, and this class includes olaparib and talazoparib. Type III PARPi have a pro-release effect from the DNA break and include rucaparib, niraparib and veliparib. Type III PARPi decrease HD dynamics and favor the closed conformation of the HD that does not contribute to DNA binding (*33*). The conversion of the type III PARPi veliparib into type I PARPi UKTT15 resulted in a compound that was more efficient at killing cancer cells, suggesting that reverse allostery can contribute to PARPi potency.

In the current study, we have analyzed the ability of PARPi to induce reverse allostery in PARP2. Recent work has shown that niraparib, talazoparib, and to a lesser extent olaparib are able to restrict the mobility of GFP-labeled PARP2 molecules within PARP2 foci that formed at sites of DNA damage in cells (*34*). This result is different from what was observed for PARP1, where none of the tested inhibitors could elicit a reduction in PARP1 mobility within foci (*35*). Instead, a rapid exchange of PARP1 was observed even in the presence of persistent foci induced by PARPi. Consistent with what was observed in cells, we show that PARPi have completely different effects on PARP1 compared to PARP2 in terms of reverse allostery. Indeed, niraparib, talazoparib, rucaparib, and to a lesser extent olaparib all act as type I inhibitors in PARP2, increasing DNA binding affinity and retention on DNA. In contrast, veliparib is the only PARPi behaving similarly between PARP1 and PARP2 as a type III inhibitor. Comparison of structures of PARP1 and PARP2 in complex with DNA suggests that differences in helix F positioning in the HD can explain the specific effect of PARPi on the two proteins. We show that a mutation on helix F can partly mimic the effect of some PARPi in our biochemical experiments and in cells. We also tested the new PARP1-selective compound AZD5305 (*36, 37*) for reverse allostery in PARP1 and PARP2 and observed that this compound exerts a clear a pro-retention effect on PARP2. Overall, our results indicate that PARPi have drastically different outcomes on DNA retention in PARP1 and PARP2 due to reverse allostery and suggest that some of these effects contribute to PARP2 trapping in cells. This PARP2-specific behavior could contribute to the side-effects that are observed with current PARPi that have been attributed to PARP2, and our molecular understanding of the process suggests ways that inhibitors could be designed to avoid or enhance these allosteric effects.

## Results

### Clinical PARPi induce PARP2-specific retention on DNA breaks

Having established a classification of PARPi based on their ability to affect DNA break retention in PARP1 (*33*) (Fig. 1), we wanted to test the PARPi effect on PARP2 ability to bind and persist on DNA breaks. We first used a fluorescence polarization (FP) assay where PARP2 binds to a fluorescent DNA dumbbell probe containing a central 5’ phosphorylated nick. An unlabelled DNA is then added as a competitor and the decrease in FP is monitored over time. Niraparib, talazoparib, and rucaparib all strongly increased retention of PARP2 on the DNA break (about 20-fold) while olaparib had a more intermediate effect (about 5-fold) (Fig. 2A,C). EB47, a control compound that mimics NAD^+^ and that is not used in the clinic, also increased retention of PARP2 on DNA. In contrast, veliparib had no effect (Fig. 2A, C). The effect of PARPi were similar whether or not HPF1 was present in the reaction (fig. s1). The PARPi effects on PARP2 were quite different from previous observations with PARP1, where niraparib and rucaparib were classified as type III inhibitors, increasing release from the DNA break, and talazoparib and olaparib were classified as type II inhibitors, having mild to no impact on DNA retention (*33*). Since the experiments that established the PARPi classification for PARP1 were performed using a DNA nick that did not carry a 5’P, we tested the reverse allosteric effect of PARPi on PARP1 using the same probe as for PARP2 to see if the 5’P would affect the results. PARP1 is activated similarly by these two types of DNA damage, while PARP2 shows a strong preference for 5’P breaks (*38*). The PARPi reverse allosteric effect observed for PARP1 did not change whether or not the DNA break carried a 5’P (Fig, 2B, D, fig. s2). These results clearly show that PARPi have different impacts on reverse allostery in PARP1 and PARP2.

**Fig. 2.**
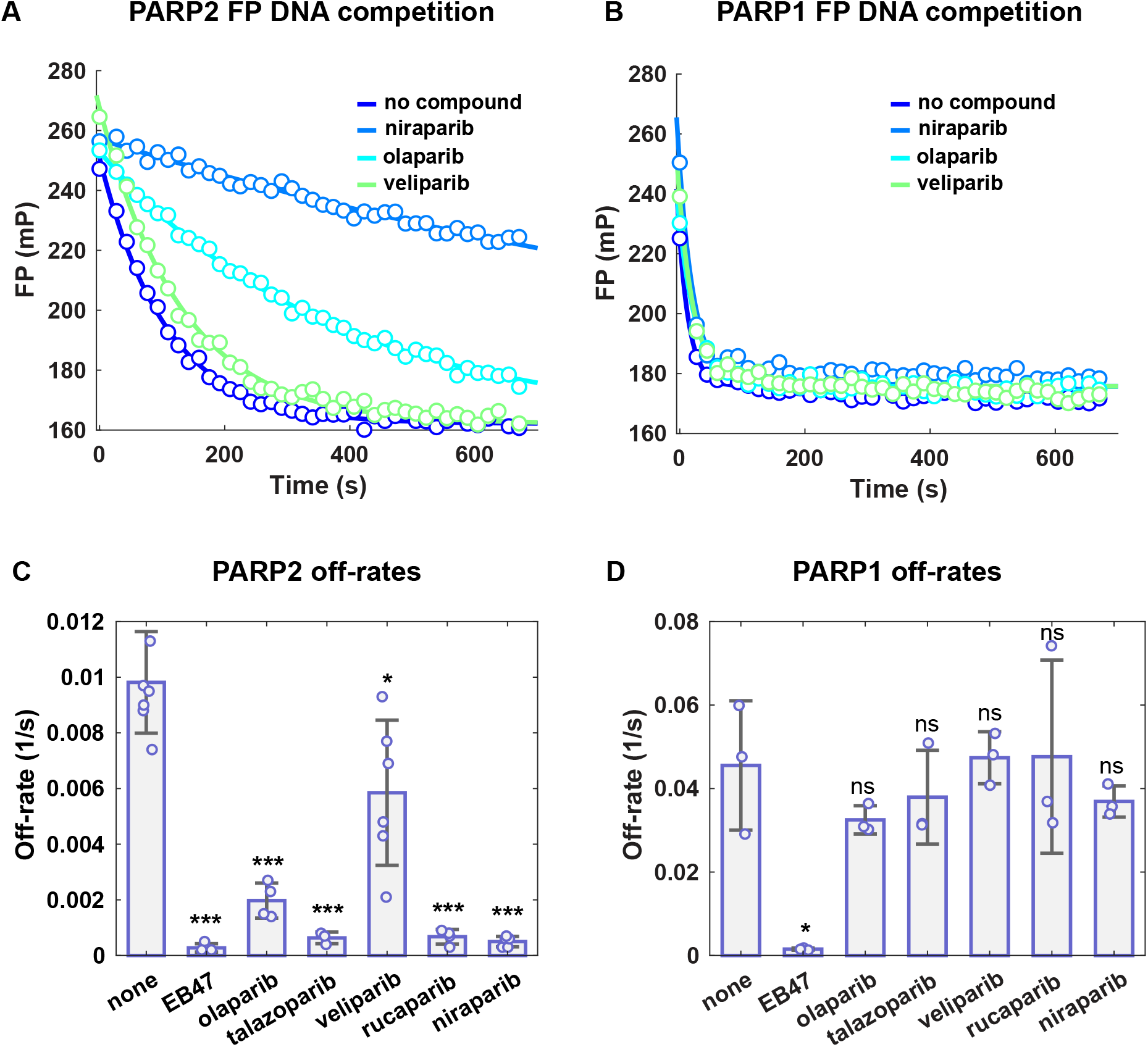
Fluorescence polarization DNA competition experiments comparing the effect of PARPi on reverse allostery in PARP2 and PARP1. (**A**) PARP2 (40 nM) was incubated with a dumbbell DNA probe containing a central 5’P nick (5 nM) for 30 minutes at room temperature in the presence of DMSO or inhibitors (100 µM). A competitor unlabelled DNA (2 µM) was added and FP was measured over time. A single exponential was fit to the data in Matlab to obtain an off-rate. (**B**) Same analysis as in **A**, but using PARP1. (**C**) Off-rates shown for PARP2 are an average of 3 to 6 independent experiments performed as in **A**. Each bar represents the mean value and the error bar corresponds to the standard deviation. The points represent the off-rate value for each individual experiment. (**D**) Same analysis as in **C** but using PARP1. Two-sample two-sided *t*-tests were used to compare the off-rate values between PARPi treatments and control samples with DMSO (none). * asterisks indicate *P* < 0.05, ** asterisks indicate *P* < 0.005, *** asterisks indicate *P* < 0.0005, and ns indicates not significant.

We next used surface plasmon resonance (SPR) to determine how PARPi affect the DNA binding kinetics of PARP2 (Fig. 3A-C), with the additional advantage that SPR is better suited than the FP assay to distinguish between type II and type III inhibitors. Indeed in PARP1, veliparib, rucaparib and niraparib did not show type III behaviors in the FP assay (Fig. 2D, fig. s2), in contrast to what was observed by SPR (*33*). A biotinylated DNA dumbbell carrying a 5’P nick was immobilized on a streptavidin coated chip. PARP2 was flowed at various concentrations in the presence of PARPi or DMSO. The association constant increased slightly in the presence of all PARPi compared to the control with DMSO (Fig. 3A). In contrast, the dissociation constant and the K_D_ were decreased substantially in the presence of talazoparib, rucaparib, niraparib, and to a lesser extent olaparib (Fig. 3B, C, fig. s3). Table 1 summarizes the affinities obtained by SPR for PARP2 for the 5’P nick DNA in the presence of the various inhibitors. The increase in affinity due to EB47, talazoparib, rucaparib, and niraparib is about 6- to 8-fold, and 2.5-fold for olaparib. In contrast, veliparib slightly decreased the affinity of PARP2 for DNA (1.2-fold).

**Table 1.** PARP2 DNA binding affinity measurements by SPR.

| PARP2 | Average $K_D$ (nM) |
| --- | --- |
| WT | 6.76 ± 0.29 |
| WT EB47 | 0.94 ± 0.21 |
| WT olaparib | 2.76 ± 0.49 |
| WT talazoparib | 0.83 ± 0.08 |
| WT veliparib | 8.09 ± 0.59 |
| WT rucaparib | 0.81 ± 0.09 |
| WT niraparib | 1.31 ± 0.02 |
| I318A | 1.60 ± 0.11 |
| I318A niraparib | 0.63 ± 0.12 |
| I318A veliparib | 1.68 ± 0.31 |
| I318V | 3.06 ± 0.18 |
| I318V niraparib | 1.31 ± 0.08 |
| I318V veliparib | 4.73 ± 0.50 |

**Fig. 3.**
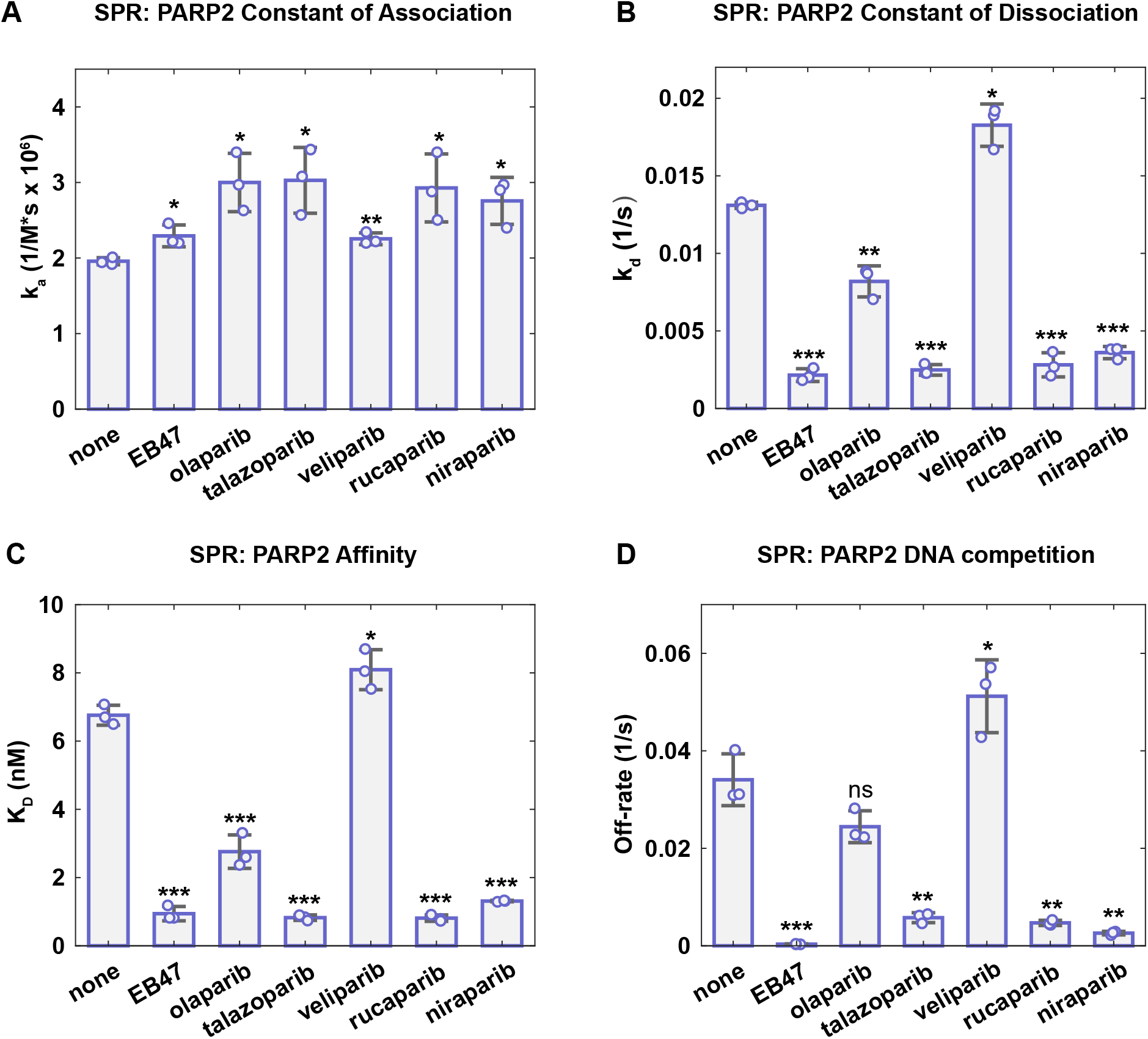
SPR experiments showing the effect of PARPi on reverse allostery in PARP2. (**A**-**C**) A streptavidin coated chip was used to capture a biotinylated dumbbell DNA containing a central 5’P nick (20 to 40 nM). PARP2 was flowed on the chip at various concentrations in the presence of DMSO or PARPi (5 µM). A 1:1 binding model was fit to the data in TraceDrawer (Reichert) to yield an association constant (k_a_) (**A**), a dissociation constant (k_d_) (**B**), and equilibrium dissociation constant (K_D_: k_a_/ k_d_) (**C**). The bars represent an average of 3 independent experiments and the error bars represent the associated standard deviations. The points represent the value obtained for each individual experiment. (**D**) PARP2 was flowed at 60 nM on a streptavidin coated chip coupled to biotinylated DNA in the presence of DMSO or PARPi (5 µM). At the time of dissociation, an external valve was used to inject non-biotinylated competitor DNA with DMSO or PARPi (5 µM). Off-rates were calculated in Matlab using a single exponential. The bars represent the average of 3 independent experiments and the error bars represent the associated standard deviations. The points represent the values obtained for each individual experiment. Two-sample two-sided *t*-tests were used to compare the k_a_, k_d_, and K_D_ values between samples with PARPi and samples with DMSO. * asterisks indicate *P* < 0.05, ** asterisks indicate *P* < 0.005, *** asterisks indicate *P* < 0.0005, and ns indicates not significant.

We next used SPR to measure the off-rates of PARP2 from the 5’P nick biotinylated DNA in the presence of a DNA competitor, similar to the FP competition assay. PARP2 was flowed on the DNA coated chip in the presence of DMSO or inhibitor. During the dissociation phase, a competitor 5’P nick DNA was added with DMSO or inhibitor and the release of PARP2 from DNA was measured over time (Fig. 3D, fig. s4). PARPi affected PARP2 release from DNA similarly to what was observed using the FP competition experiments. However, in this case, veliparib showed a pro-release effect that was not observed in the FP assay, but that is consistent with the observed slight decrease in affinity in the SPR kinetics experiment (Fig. 3A-C). Overall, the FP and SPR experiments clearly indicate that most PARPi do not fall in the same reverse allosteric class in PARP2 compared to PARP1. For PARP2, the type I class includes EB47, talazoparib, rucaparib, niraparib, and olaparib. The only type III PARPi was veliparib, and no type II PARPi were found for PARP2 among the ones tested.

### Determinants of the pro-retention effects in PARP2

We next tested if WGR-HD communication was important for the pro-retention reverse allosteric effect that we observe in PARP2 with some PARPi. In PARP1, the communication between the regulatory domains was shown to be important for the effect of type I inhibitor EB47 (*33*). We tested the WGR mutant N116A for its ability to respond to niraparib in the FP release assay. N116 contacts the C-terminal end of helix E and adjacent loop in the HD of PARP2 (Fig. 4A). Niraparib was not able to promote retention of the N116A mutant, suggesting that contact between the WGR and the HD are important for the pro-retention effect of niraparib on PARP2 (Fig. 4B, C). Additionally, the N116A mutant was competed off the DNA probe much more rapidly than PARP2 WT, consistent with the DNA binding deficiency of the N116A mutant (*16*). This result is coherent with the finding in PARP1 that the HD contributes to DNA binding by interacting with the WGR domain (*22*).

**Fig. 4.**
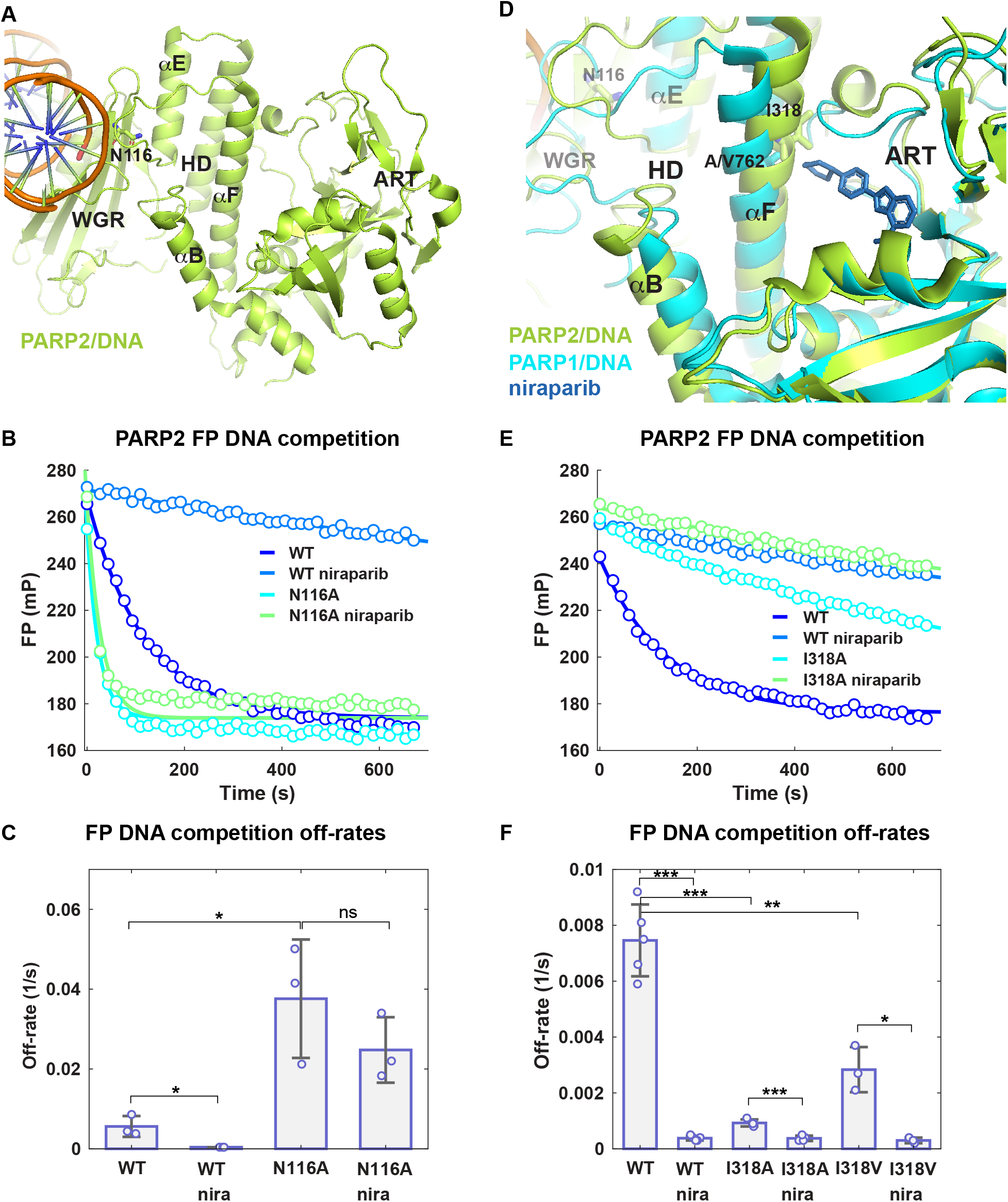
PARP2 mutants alter PARPi reverse allosteric effect. (**A**) The cryo-EM structure of PARP2/HPF1/DNA complex (6X0L) is shown. HPF1 has been omitted for clarity. The location of WGR residue N116 is indicated. (**B**) Fluorescence polarization DNA competition experiment with PARP2 WT and mutant N116A with or without niraparib (100 µM). (**C**) Off-rates were determined by fitting a single exponential to the data in **B**. Averages of 3 independent experiments are shown and the error bars represent the associated standard deviations. The points represent the value obtained for each individual experiment. (**D**) The crystal structure of PARP1/DNA complex (4DQY) and the crystal structure of PARP1 CAT bound to niraparib (4R6E, only niraparib shown) were aligned to the cryo-EM structure of PARP2/HPF1/DNA complex (6X0L, HPF1 was omitted for clarity). The position of I318 in PARP2 and V/A762 in PARP1 are shown. (**E-F**) Same as in **C** and **D** for I318A and I 318V mutants. Two-sample two-sided *t*-tests were used to compare the off-rates between samples as indicated. * asterisks indicate *P* < 0.05, ** asterisks indicate *P* < 0.005, *** asterisks indicate *P* < 0.0005, and ns indicates not significant.

In order to understand the differential effects of PARPi in PARP1 and PARP2, we aligned the crystal structure 4DQY of PARP1/DNA (*12*), the cryo-EM structure 6X0L of PARP2/DNA (*18*), and the crystal structure 4R6E of niraparib bound to PARP1 CAT (*39*)(Fig. 4D). The alignment was performed by superposing the ART domains. Niraparib comes close to the N-terminal part of helix F in a region where the structures of PARP1 and PARP2 differ in terms of the positioning of helix F relative to the ART. Structural alignments of crystal structures of PARP1 or PARP2 CATs with other compounds show that rucaparib and talazoparib also come close to the N-terminal section of helix F (fig. s5). In contrast, olaparib approaches the middle part of helix F, and EB47 the C-terminal part. The smaller veliparib compound is the farthest from helix F.

In PARP1, the N-terminal part of helix F is bent away from the ART, providing more space for the niraparib compound to fit (Fig. 4D). In contrast, in PARP2 the helix is straighter and appears to be in a position that would clash with niraparib. Therefore, niraparib might provide some distortion at the N-terminus of helix F in PARP2, pushing the HD into a conformation that favors interaction with the WGR domain and increases DNA binding and retention. Sequence conservation analysis of PARP1 and PARP2 in this region indicated that I318 in PARP2 is equivalent to alanine/valine at position 762 in PARP1, the site of a natural variant in PARP1 (*40*). We reasoned that mutating I318 to an alanine or a valine in PARP2 might provide more space for niraparib to fit and consequently abrogate its pro-retention effect. The I318A and I318V mutants were created and tested in the FP release assay (Fig. 4D-F). Surprisingly, the I318A mutation alone increased DNA retention even in the absence of niraparib by about 8-fold (Fig. 4E, 4F). The I318V mutation also increased DNA retention but had an intermediate effect of about 2.5-fold (Fig. 4F). Both mutants still exhibited a decrease in off-rate from DNA when niraparib was added, although to a smaller extent than observed with PARP2 WT. SPR was used to determine the DNA binding affinities of the two mutants and showed a decrease in K_D_, representing an affinity increase of about 4-fold for I318A and 2-fold for I318V, consistent with the FP results (Table 1, fig. s6). Therefore, rather than disrupting the effect of niraparib, the I318A and I318V mutations have partly mimicked its effect on PARP2, increasing DNA binding affinity and retention. We expect that these mutations alter the conformation of the N-terminus of helix F in a way that favors the open HD conformation that interacts with the WGR domain and increases DNA binding affinity. Niraparib binding is likely to displace the N-terminal region of helix F and therefore have similar consequences on PARP2 DNA retention. Talazoparib and rucaparib, which occupy a similar space, would have a similar effect as niraparib, as observed in our biochemical experiments.

### PARP2 I318A accumulates at higher levels than PARP2 WT at DNA damage sites in cells

Recent live cell imaging experiments demonstrated that PARP2 mobility at sites of DNA damage was reduced by the PARPi niraparib, talazoparib, and to a lesser extent olaparib (*34*). It was hypothesized that reverse allostery was playing a role in the trapping exerted by these compounds. These results contrast with what was observed for PARP1 where none of the PARPi tested could induce similar reduction in PARP1 mobility at damage sites (*35*). Instead, PARP1 was shown to rapidly exchange at sites of damage. In order to test if reverse allostery is involved in inducing trapping of PARP2 at DNA damage sites, we used our I318A mutant that partly mimicked the effect of niraparib treatment. PARP1/2 double knock-out TERT-immortalized human retinal pigment epithelial-1 (RPE-1) cells were transfected with GFP-PARP2 WT or I318A. XRCC1 binds to PAR, so RFP-XRCC1 was used to monitor PAR formation. After DNA damage was induced by laser micro-irradiation in a defined nuclear region, foci formation and persistence were monitored over time in live cells (Fig. 5). Consistent with our previous work (*34*), PARP2 WT localized to sites of DNA damage immediately after micro irradiation (within 2 seconds), peaked within 60 seconds, and persisted over the time-course tested. The PARP2 I318A mutation promoted localization to sites of damage (Fig. 5A, B, C), consistent with our biochemical results that indicated increased retention on a DNA break. Interestingly, PARP2 I318A stimulated recruitment of XRCC1 to DNA breaks as observed by an increase in foci intensity compared to PARP2 WT (Fig. 5A, D, E). This observation is consistent with the fact that the I318A mutant showed increased DNA-independent activity compared to PARP2 WT, in particular in the presence of HPF1 (fig. s7). This increase is likely due to the fact that the mutations favor the HD conformation that engages the WGR and therefore partially relieves the steric blockage of the autoinhibitory HD and opens the ART for NAD^+^ binding (*20, 21*). Additionally, since the HD in its closed conformation impedes HPF1 binding, the opening of the HD in the I318A mutant could increase the ability of HPF1 to bind PARP2 and stimulate initiation activity as described previously (*8*). The elevated catalytic activity of the I318A mutant could therefore lead to increased recruitment of PAR binding factors such as XRCC1.

**Fig. 5.**
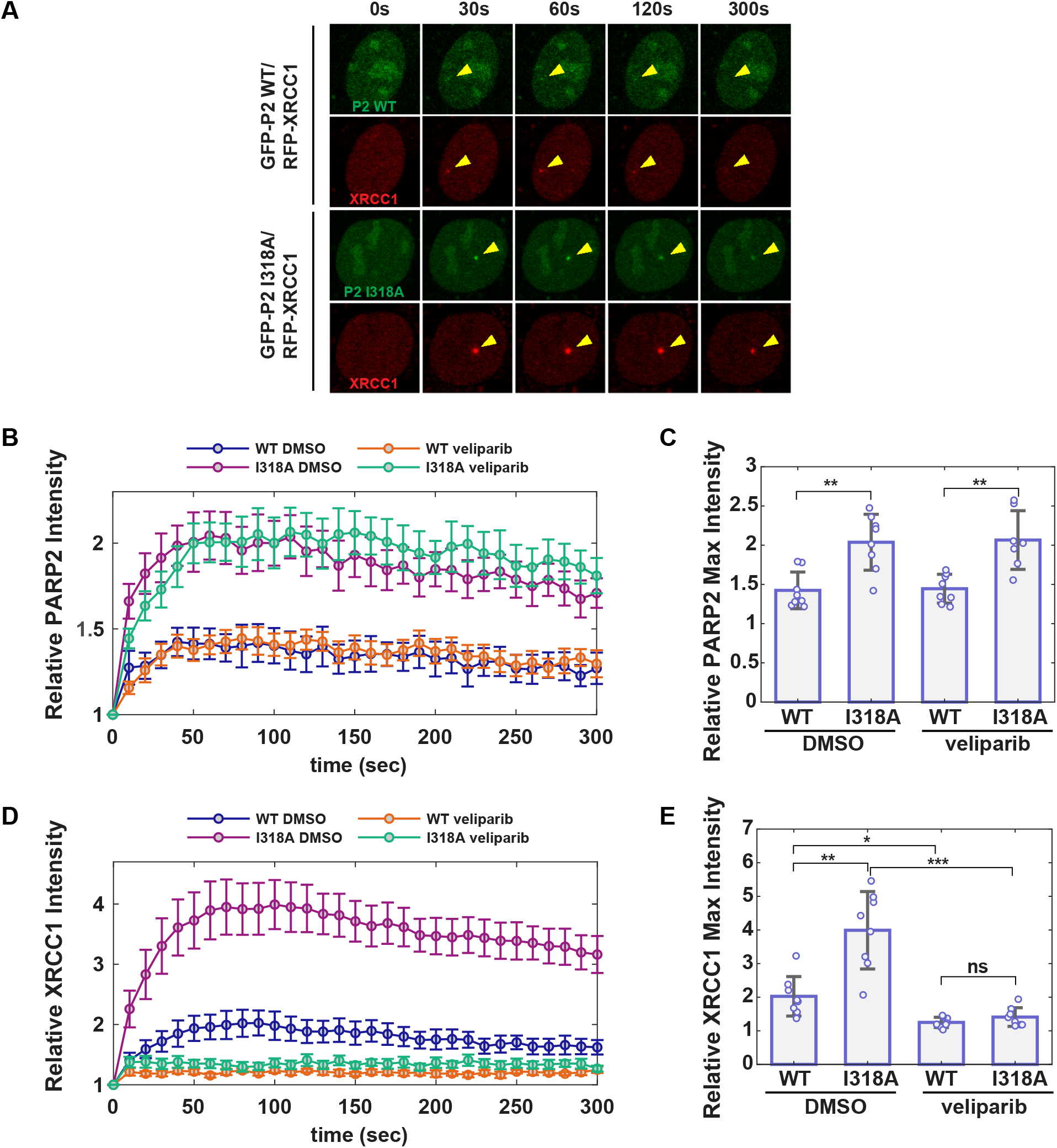
PARP2 I318A accumulates more efficiently than PARP2 WT at DNA damage sites in cells. (**A**) Representative images of laser-induced GFP-PARP2 WT or I318A mutant foci, and RFP-XRCC1 foci in PARP1/2 KO RPE-1 cells. The arrowheads point to the site of micro-irradiation. (**B**) The relative intensity of PARP2 at DNA damage sites (normalized to the intensity before irradiation) in the presence of DMSO or veliparib. The points and error bars represent the averages and standard errors, respectively. (**C**) The maximum relative intensity of GFP-PARP2 from **B**. The bars represent the average of the maximum relative intensity obtained for the cells in one representative experiment out of 3 consistent biological repeats. Each point shown represents the maximum relative intensity obtained for one out of 7 to 8 cells. The error bars correspond to the standard deviations. Two-sample two-sided *t*-tests were used to compare the relative PARP2 foci intensity between samples as indicated. * asterisks indicate *P* < 0.05, ** asterisks indicate *P* < 0.005, *** asterisks indicate *P* < 0.0005, and ns indicates not significant. (**D**) and (**E**) are the same analysis as in (**B**) and (**C**) but using XRCC1.

Since PAR formation on PARP1 and PARP2 attenuates their DNA binding, we added the PARPi veliparib to the cells in order to remove the contribution of catalytic activity in the kinetic experiments, and to thus isolate the DNA binding activity (Fig. 5B, C, D, E, fig. s8). For both PARP2 WT and the I318A mutant, the addition of veliparib had no effect on the accumulation of PARP2 to sites of damage, and the I318A mutant still showed higher foci intensity. XRCC1 recruitment was diminished in both cases, indicating that veliparib indeed inhibits catalytic activity under these conditions. We interpret this result to indicate that in the absence of catalytic activity, the mutant accumulated at higher levels at the sites of damage, consistent with our biochemical results showing an increase in DNA binding affinity and retention for I318A. We next added niraparib to PARP2 WT and I318A cells and observed that niraparib has a larger impact on recruitment and retention to the DNA break than the I318A mutation by itself without PARPi (Fig. s8). This result is consistent with our biochemical data showing that the I318A mutant does not fully recapitulate the reverse allosteric effect of niraparib on PARP2 DNA binding and retention. These experiments show that the increase in reverse allosteric DNA retention observed *in vitro* for the I318A mutant translates into an increase in accumulation at sites of damage in cells, supporting the idea that reverse allostery contributes to niraparib-mediated trapping of PARP2 in cells.

### The AZD5305 compound induces DNA break retention in PARP2

AZD5305 is a novel PARPi that has been developed as a selective PARP1 inhibitor, in an attempt to decrease the hematological toxicity associated with the clinical PARPi that has been attributed to targeting other PARP family members and in particular PARP2 (*36, 37*). AZD5305 was reported to have a 460-fold selectivity for PARP1 over PARP2 with an IC_50_ of 0.003 µM in PARP1 compared to 1.4 µM in PARP2 (*36*), and a similar selectivity for inhibition of PARP1 over PARP2 in cells (*37*). Aligning the crystal structures of PARP1 CAT bound to an analog of AZD5305 (compound 22, (*36*)), PARP1 CAT bound to niraparib, and PARP2 CAT bound to olaparib showed that AZD5305 contacts the HD in the middle part of helix F similar to olaparib (Fig. 6A). We first tested the effect of AZD5305 on PARP1 retention on DNA breaks by FP (Fig. s9). Under these conditions, AZD5305 had no measurable effect on the off-rate measured for PARP1 by FP compared to the DMSO control. In contrast, AZD5305 increased PARP2 retention on a DNA break with an intermediate effect, similar to olaparib at 100 µM concentration (Fig.6B, Fig. s10). We tested the off-rates of PARP2 from DNA by FP in the presence of various concentrations of AZD5305 and olaparib (Fig. 6C, D). AZD5305 showed a decrease in PARP2 off-rate starting at 1 µM, while the effect of olaparib was already visible at 0.1 µM. Therefore, despite its high selectivity for PARP1, AZD5305 could still have an effect in cell experiments due to reverse allostery targeting PARP2, depending on the concentrations used.

**Fig. 6.**
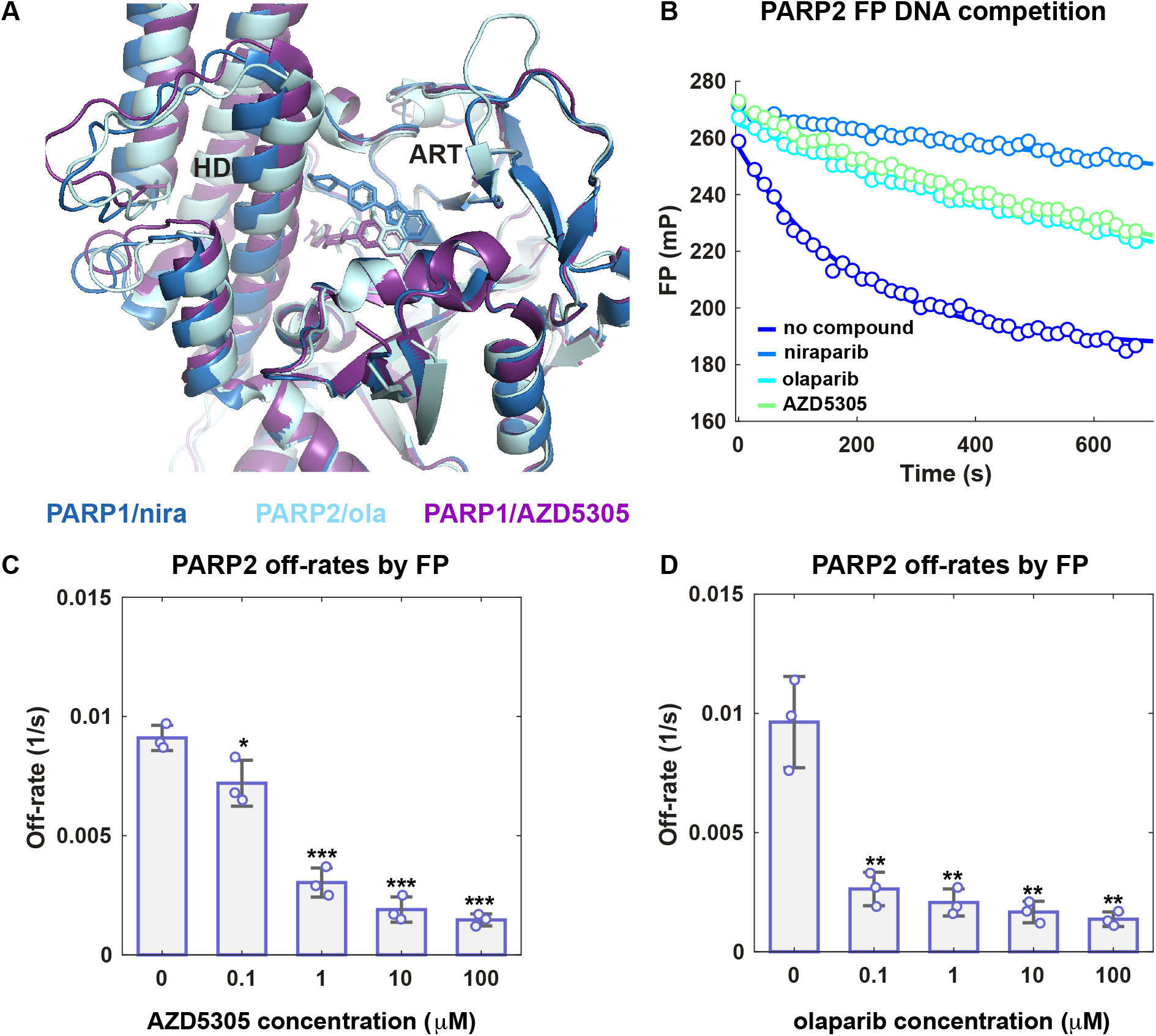
Effect of AZD5305 on PARP2 retention on DNA. (**A**) The crystal structure of PARP1 CAT bound to niraparib (4R6E) was aligned to the crystal structure of PARP2 CAT bound to olaparib (4TVJ) and the crystal structure of PARP1 CAT bound to an analog of AZD5305 (7ONT). (**B**) Fluorescence polarization DNA competition experiment with PARP2 WT in the presence of niraparib, olaparib or AZD5305 (100 µM). (**C-D**) Off-rates were determined by fitting a single exponential to the fluorescence polarization DNA competition data at various concentrations of AZD5305 (**C**) or olaparib (**D**). The bars represent averages of 3 independent experiments and the error bars represent the associated standard deviations. The points represent the value obtained for each individual experiment. Two-sample two-sided *t*-tests were used to compare the relative off-rates between samples with PARPi and samples with DMSO. * asterisks indicate *P* < 0.05, ** asterisks indicate *P* < 0.005, *** asterisks indicate *P* < 0.0005.

## Discussion

In contrast to what was proposed earlier (*26, 41*), the analysis of PARPi effect on PARP1 demonstrated that none of the clinical inhibitors have a pro-retention, reverse allosteric effect that enhances PARP1 binding to DNA (*33*). In fact, some of the clinical PARPi (niraparib, rucaparib and veliparib) have the opposite effect, increasing PARP1 release from DNA damage. Here we have shown that in PARP2, a reverse allosteric pro-retention effect is observed for several clinical PARPi, namely talazoparib, niraparib, rucaparib, and to a lesser extent olaparib. This difference between PARP1 and PARP2 could be explained by the positioning of HD helix F, which appears to act as a sensor for compounds bound to the catalytic site. Indeed, in PARP2, the first half of helix F is bent toward the ART compared to PARP1. Consequently, PARPi that contact or come close to this section of the helix when bound to the active site seem to induce an increase in DNA retention in PARP2, which is not observed for PARP1. We have shown previously that the HD domain contributes to the ability of PARP1 to bind to DNA (*22*). In the closed conformation, when PARP1 is not bound to DNA, the HD is fully bound to the ART domain and has limited contact with WGR. When PARP1 binds to DNA, the HD becomes more mobile with dramatic changes in helix dynamics (*20*), and samples a conformation where it interacts with WGR, which leads to an increase in PARP1 affinity for DNA (*22*). In this conformation, the ART is open for NAD^+^ binding (*21*). When NAD^+^ or type I inhibitors bind to the catalytic site, they stabilize the open conformation of the HD and therefore increase DNA binding (*21, 33*).

In PARP2, type I inhibitors that contact the N-terminal part of helix F (niraparib, talazoparib and rucaparib) could have a similar effect by disturbing the local structure of helix F and favoring the HD conformation that interacts with the WGR domain. Mutation of residue I318 located at the N-terminal part of helix F seems to mimic this pro-retention effect, likely by inducing a similar structural perturbation as a type I inhibitor. Interestingly, the I318A mutant also has increased DNA-independent activity *in vitro* and shows overactivity in cells, consistent with the idea that the HD conformation is altered and the ART is partially opened for NAD^+^ binding, even in the absence of DNA. In contrast to niraparib, talazoparib, and rucaparib, the PARPi olaparib contacts the middle portion of helix F, where the helix adopts a more similar position in PARP1 and PARP2. However, in PARP2 a larger Glu residue (Glu335) has replaced the Asp residue present in PARP1 (D766). This difference could explain why olaparib has modest type I behavior in PARP2 but type II behavior in PARP1, where there is more space to accommodate the compound and therefore no clash with helix F. The non-clinical PARPi EB47 has a similar pro-retention effect in PARP1 and PARP2 and contacts the C-terminal part of helix F. In this case, EB47 could displace this section of the helix and still favor the HD-WGR interaction that increases DNA binding affinity. Veliparib, is the only PARPi that shows mild pro-release activity in PARP2. Veliparib is the smallest compound and does not come as close to helix F as other PARPi. The veliparib pro-release effect could be due to stabilizing ART contacts that in turn lead to rigidification of the HD, which would then be less likely to bind to WGR. That kind of rigidification or decrease in HD dynamics was observed in PARP1 upon binding to veliparib, niraparib and rucaparib, which all act as type III inhibitors in PARP1 (*33*).

A recent crystal structure of PARP2 has shown a large displacement of the ART away from the HD and the WGR when PARP2 binds to DNA (*17*). This change in conformation separating the ART from the HD was not observed in the cryo-EM structure of PARP2/ HPF1/nucleosomes (*18*), or in PARP1/DNA crystal structures (*12, 22*). The mechanism is not clear for how a PARPi bound ART that is displaced from the HD could influence WGR binding to DNA. However, it is possible that type I PARPi in PARP2 actually promote this displacement and prevent rebinding of the ART to the HD, and therefore favor the fully open conformation of the HD that interacts with the WGR and increases DNA binding. The crystal structure of a mutant of PARP1 that favors the active state bound to DNA has shown a rotation of the ART that opens the NAD^+^ binding site (*22*), and this arrangement had not been captured in the crystal structure of PARP1 WT bound to DNA (*12*). These recent structures (*17, 22*) exemplify the mobility of the ART in both PARP1 and PARP2, which is necessary to free the HD and allow it to fully bind WGR and contribute to DNA binding.

There has been a recent push to develop PARP1-specific PARPi that would not target other PARP family members and in particular PARP2. Indeed, the hematological toxicity of current clinical PARPi that leads to side-effects has been attributed mostly to targeting of PARP2 (*28, 37*). Our study suggests that some of the side-effects observed could be due to the reverse allosteric retention of PARP2 on DNA induced by type I inhibitors such as talazoparib, niraparib, rucaparib, and olaparib. In the case of AZD5305, we observed clear type I behavior in PARP2 but not in PARP1. However, the weak affinity of AZD5305 for PARP2 is likely to prevent its binding to PARP2 in cells, which is consistent with the lower hematological toxicity observed in mice for this compound compared to other PARP2 type I clinical inhibitors (*37*). We have shown recently that reverse allostery can play a role in modulating the ability of a PARPi to kill cancer cells by converting a type III PARPi (veliparib) into a type I inhibitor (UKTT15) (*33*). However, inhibitory potency and binding kinetics also play a critical role in determining the overall efficiency of a PARPi (*31, 32*). Since none of the current clinical PARPi have a pro-retention reverse allosteric effect on PARP1, it would be interesting to design and study novel PARPi that do have this characteristic for PARP1 but that do not target PARP2. This direction could lead to the discovery of potentially more potent PARPi that combine reverse allostery and catalytic inhibition to trap PARP1 on DNA with potentially limited side-effects.

## Material and methods

### Expression constructs and mutagenesis

PARP2 (isoform 2, residues 1 to 570) and PARP1 (residues 1 to 1014) were expressed from a pET28 vector with an N-terminal hexahistidine tag. The human HPF1 gene was synthesized for expression from a pET28 vector with an N-terminal His-tag and sumo-like tag (SMT). Site-directed mutagenesis on PARP2 was performed using the QuikChange protocol (Stratagene) and verified by automated Sanger sequencing. The DsRed-mono-C1-XRCC1 and pEGFP-C1-PARP2 plasmids were generously provided by Dr Li Lan at Massachusetts General Hospital (*42*) and Dr Xiaochun Yu at Westlake University (*43*), respectively.

### Cell lines and cell culture

PARP1/2 double knockout (DKO) RPE-1 cells were generously provided by Dr Keith W. Caldecott at the University of Sussex (*44*) and cultured in DMEM medium (GIBCO, Cat. 12430062) supplemented with 10% fetal bovine serum (FBS), MEM non-essential amino acids (GIBCO, Cat. 11140050), 2 mM glutamine, 1 mM sodium pyruvate and 50 U/ml penicillin/streptomycin (GIBCO, 15140122).

### Protein expression and purification

PARP2 WT and mutant proteins were expressed in *E. coli* Rosetta2 cells in media supplemented with 10 mM benzamide and purified as described previously using Ni^2+^-affinity, heparin affinity (250 mM to 750 mM NaCl elution gradient), and gel filtration chromatography (*38*). PARP1 (*11, 20, 45*–*47*) and HPF1 (*8*) were expressed and purified using Ni^2+^-affinity, heparin affinity, and gel filtration chromatography.

### Fluorescence polarization release assay

PARP2 WT and mutants (40 nM) were incubated with 20 nM dumbbell DNA with a central nick carrying an internal fluorescent FAM group and a 5’P (5’ P GCT GAG C/FAMT/T CTG GTG AAG CTC AGC TCG CGG CAG CTG GTG CTG CCG CGA) for 30 minutes at RT in 12 mM Hepes pH 8.0, 250 mM NaCl, 4% glycerol, 5.7 mM beta-mercaptoethanol, 0.05 mg/ml BSA in the presence of inhibitors (100 µM, or as indicated) in 1% DMSO (final concentration). Where indicated, HPF1 was present in the reaction at 5 µM. A competitor unlabelled DNA of the same sequence was added at 2 µM and FP was measured over time on a VictorV plate reader (Perkin Elmer). Experiments with PARP1 were carried out in the same conditions as for PARP2 with the exception that the reaction buffer which was 12 mM Hepes pH 8.0, 60 mM KCl, 8 mM MgCl_2_, 4% glycerol, 5.7 mM beta-mercaptoethanol, 0.05 mg/ml BSA with either the 5’P nick or the unphosphorylated nick DNA, as indicated. Off-rates were calculated in Matlab by fitting the data to a single exponential model.

### Surface plasmon resonance

SPR experiments were performed on a Reichert 4SPR biosensor instrument in the following buffer: 25 mM HEPES pH 7.4, 450 mM NaCl, 0.1 mM TCEP, 1 mM EDTA, and 0.05% Tween20. Streptavidin coated chips (Reichert) were used to capture a DNA dumbbell carrying a central 5’P nick and bearing a biotin group (20 nM to 40 nM). In the binding kinetics experiments, PARP2 was flowed on the biosensor at various concentrations as indicated (fig. s3 and s6) in the presence of DMSO or inhibitors (5 µM). All data was processed in TraceDrawer (Reichert) and double-referenced to buffer and a control channel that did not contain the immobilized DNA. The association and dissociation phases of the PARP2 titration on DNA was fit with a 1:1 binding model to obtain a constant of association *k*_a_, constant of dissociation *k*_d_, and apparent equilibrium dissociation constant K_D_ (*k*_a/_*k*_d_). In the DNA competition experiments, PARP2 was flowed over the DNA coupled chip at 60 nM in the presence of inhibitor (5 µM) or DMSO. An external valve was used to inject the competitor 5’P nick DNA (300 nM) with inhibitor (5 µM) or DMSO. Off-rates were calculated in Matlab by fitting the data to a single exponential model.

### Live-cell imaging data collection and processing

Live-cell imaging analyses were performed as previously described (*34*). Briefly, *PARP1/2 DKO* RPE-1 cells were seeded onto 35 mm diameter glass-bottom plates and transfected with plasmids encoding EGFP-PAPR2 WT or I318A, and RFP-XRCC1. Live-cell imaging was performed 24 h after transfection on a Nikon Ti Eclipse inverted microscope equipped with the A1 RMP confocal microscope system and Lu-N3 Laser Units (Nikon Inc, Tokyo, Japan). Micro-irradiation was carried out in the nucleoplasm area using a 405 nm laser (energy level ∼500 uW for a ∼0.8 m diameter region). Time-lapse images were acquired right before and after micro-irradiation with 10 s interval for a total of 5 min. The images were quantified using ImageJ (Fiji). The relative intensity of PARP2 and XRCC1 foci was calculated as the ratio of the mean intensity at each micro-irradiation damaged site to the corresponding mean intensity of the nucleus, then normalized to the intensity in the first image before micro-irradiation.

### SDS-PAGE assay

The SDS-PAGE activity assay was performed as described (*47*) using 1 µM PARP2, 1 µM DNA, HPF1 where indicated, 1 µM DNA where indicated (dumbbell with a central nick 5’ P PARP2), and 500 µM NAD^+^ for various times (see figures). SDS-PAGE loading buffer was added to the reactions prior to resolution on a 12% SDS-PAGE, which was then treated with Imperial stain for visualization.

### Statistical analysis

Data were analyzed using a Student’s t test (two-tailed). The differences were considered statistically significant at P < 0.05 (*), P<0.005 (**), and P<0.0005 (***), and n.s. indicates non-significant.

## Acknowledgments

We acknowledge support from the Canadian Institutes of Health Research (PJT173370 to JMP), the Natural Sciences and Engineering Research Council of Canada (RTI-2018-00894 to JMP), and the National Institutes of Health (R01CA259037 to JMP, and R01CA226852 and R01CA271595 to SZ).

## Author contributions

MFL performed mutagenesis, protein purification, fluorescence polarization assays, SPR experiments and activity assays. XL performed live-cell experiments. MFL and JMP wrote the manuscript with input from all coauthors. JMP and SZ directed the study.

## Data material and availability

All data are available in the main text or the supplementary materials.

