## Supplemental Data for "Clinical PARP inhibitors allosterically induce PARP2 retention on DNA"

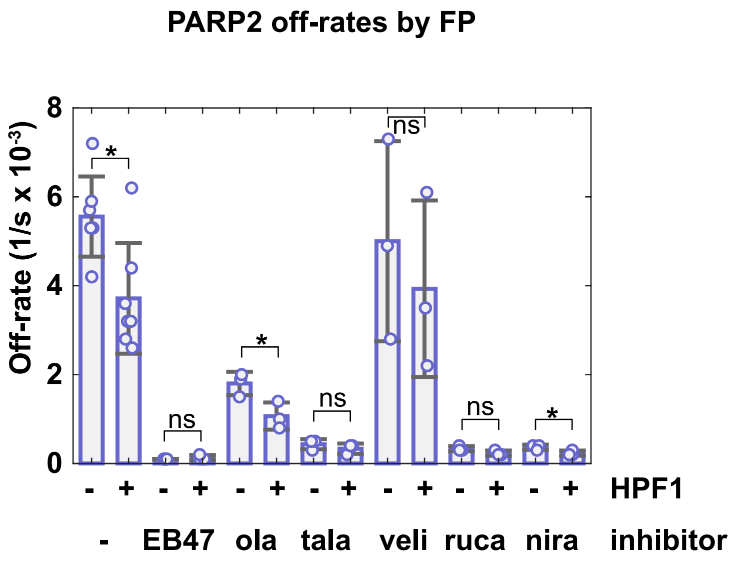


**Fig. S1. Fluorescence polarization DNA competition experiments showing the effect of PARPi on PARP2 DNA retention with and without HPF1.** PARP2 (40 nM) was incubated with a dumbbell DNA probe containing a central 5’P nick (5 nM) with or without HPF1 (5 µM) for 30 minutes at room temperature in the presence of DMSO or inhibitors (100 µM). A competitor unlabelled DNA (2 µM) was added and FP was measured over time. A single exponential was fit to the data in Matlab to obtain an off-rate. The off-rates shown are an average of 3 to 7 independent experiments. The error bar corresponds to the standard deviation. The points represent the off-rate value for each individual experiment. Two-sample two-sided *t*-tests were used to compare the off-rate values and asterisks (*) indicate $P < 0.05$. ns, indicates not significant.

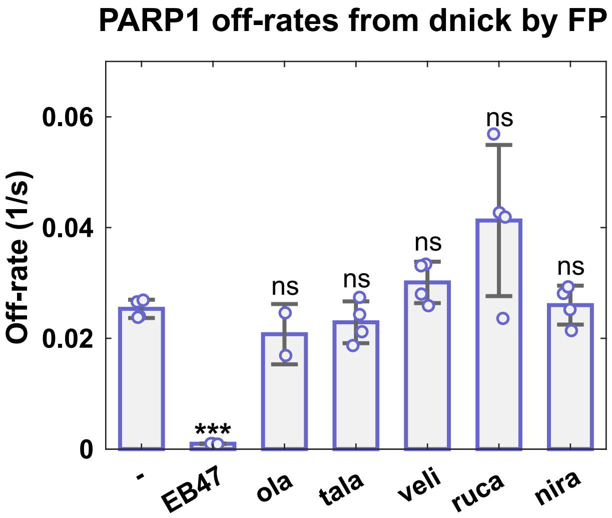


**Fig. S2. Fluorescence polarization DNA competition experiments showing the effect of PARPi on PARP1 DNA retention with an unphosphorylated nicked DNA.** PARP1 (40 nM) was incubated with a dumbbell DNA probe containing a central nonphosphorylated nick (5 nM) for 30 minutes at room temperature in the presence of DMSO or inhibitors (100 µM). A competitor unlabelled DNA (2 µM) was added and FP was measured over time. A single exponential was fit to the data in Matlab to obtain an off-rate. The off-rates shown are an average of 2 to 4 independent experiments. The error bar corresponds to the standard deviation. The points represent the off-rate value for each individual experiment. Two-sample two-sided *t*-tests were used to compare the off-rate values between samples with PARPi and samples with DMSO. Asterisks (***) indicate $P < 0.0005$. ns, indicates not significant.

### SPR binding kinetics

#### PARP2 DMSO

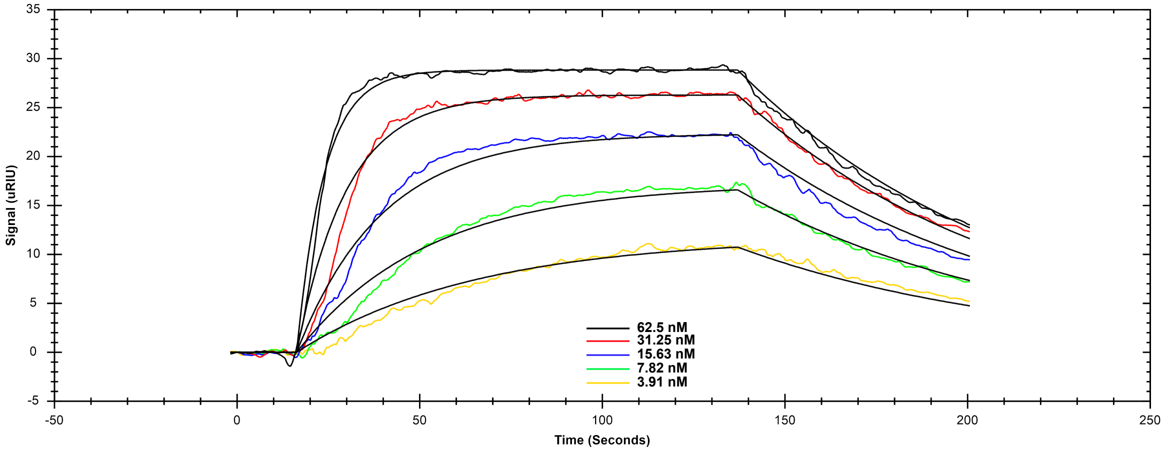


#### PARP2 EB47

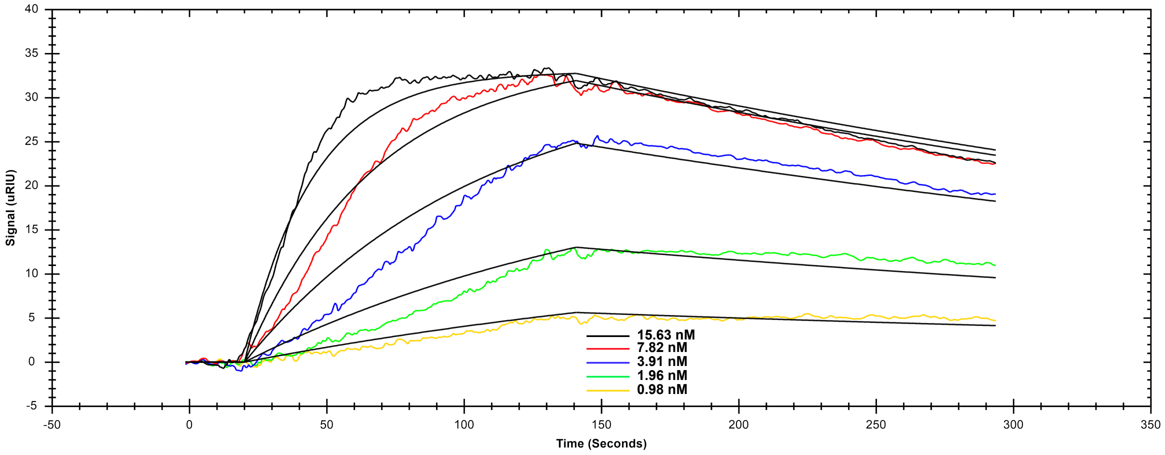


#### PARP2 olaparib

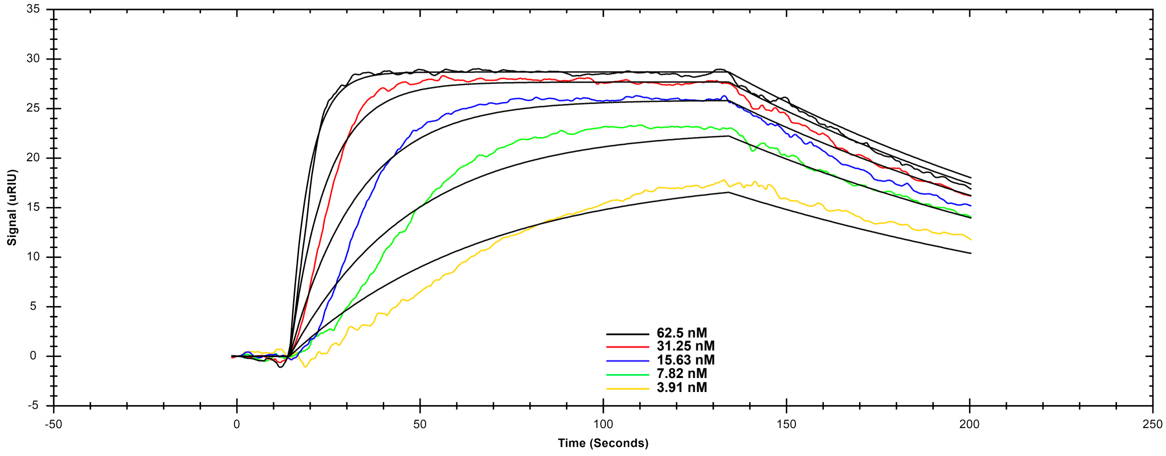

### SPR binding kinetics

#### PARP2 talazoparib

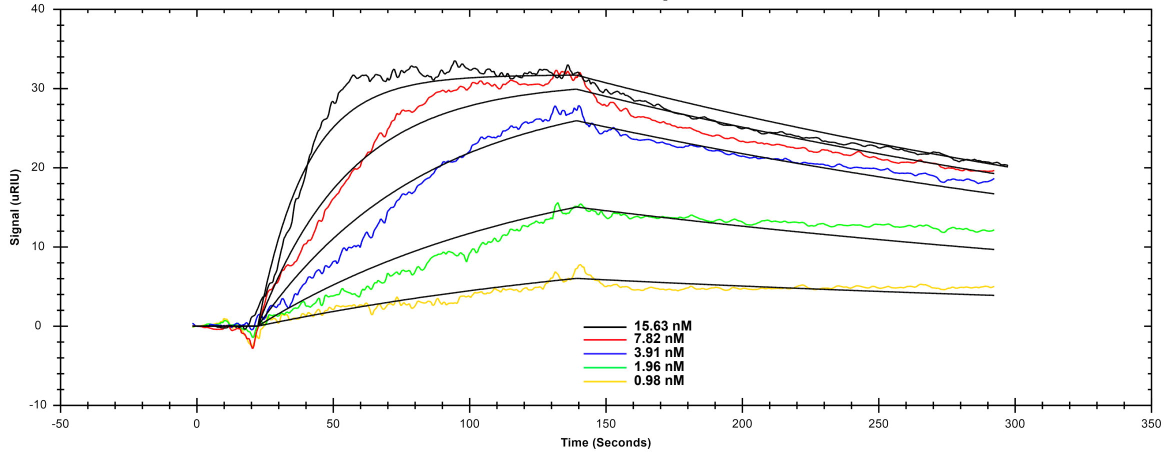


#### PARP2 veliparib

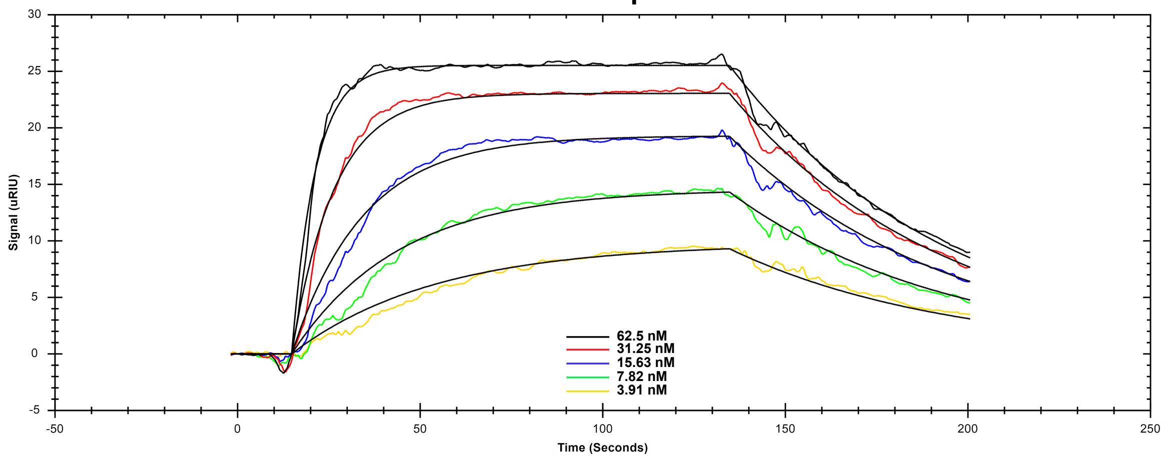


#### PARP2 rucaparib

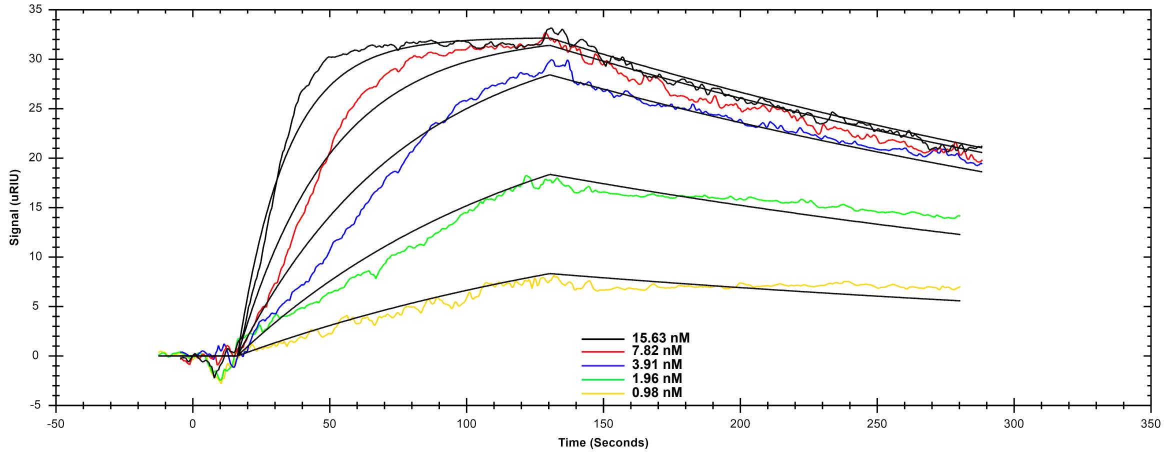

### SPR binding kinetics

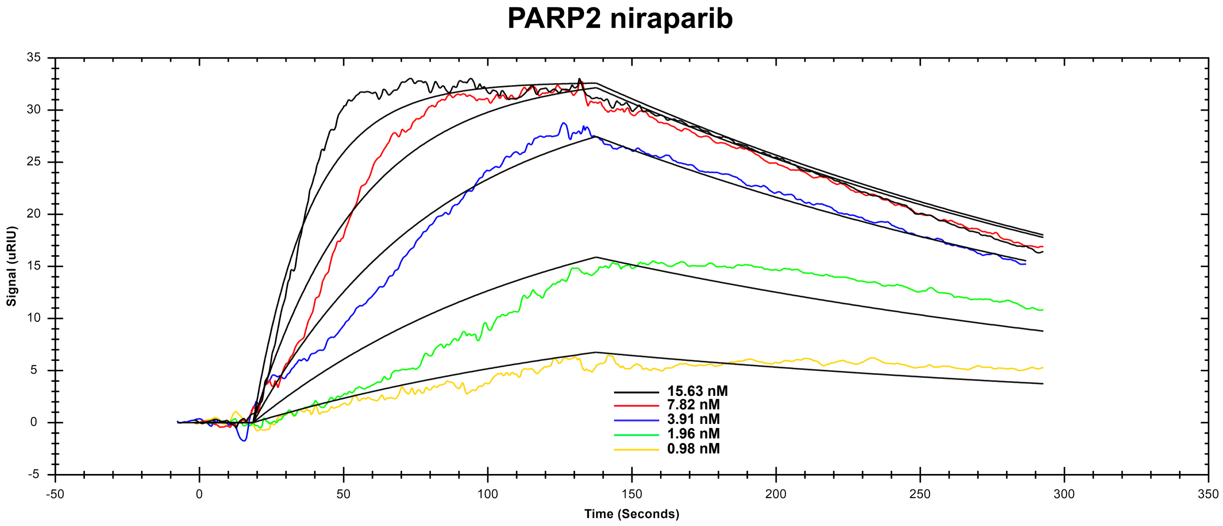


**Fig. S3. SPR binding curve titrations.** A biotinylated DNA containing a central 5'P nick was captured on a streptavidin coated chip. PARP2 WT was flowed at various concentrations over the chip as indicated on the figure in the presence of DMSO or 5 μM inhibitor. A 1:1 binding model was fit to the data in TraceDrawer (Reichert).

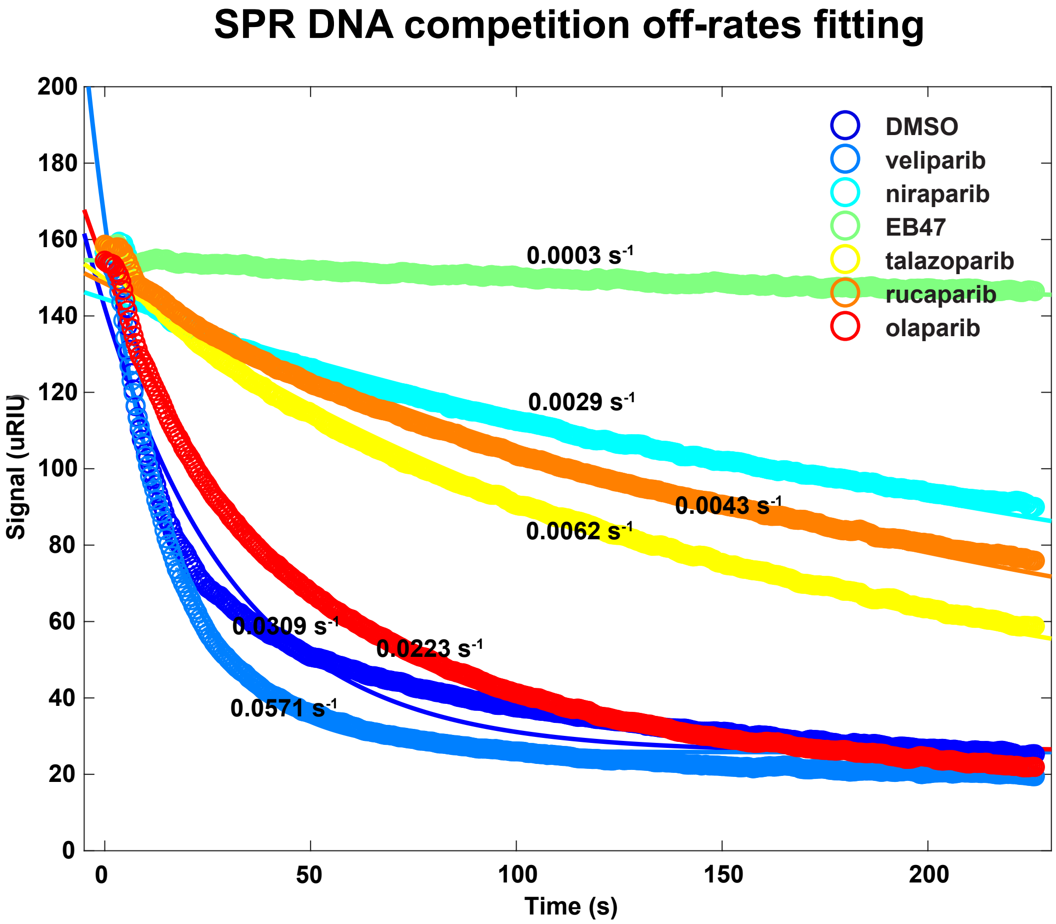


**Fig. S4. PARP2 DNA competition assay by SPR, examples of fits to the data.** PARP2 was flowed at 60 nM on the streptavidin coated chip coupled to the biotinylated DNA (5'P nick) in the presence of DMSO or inhibitors (5 μM). At the time of dissociation, an external valve was used to inject competitor unbiotinylated DNA with DMSO or inhibitor (5 μM). Off-rates were calculated in Matlab using a single exponential.

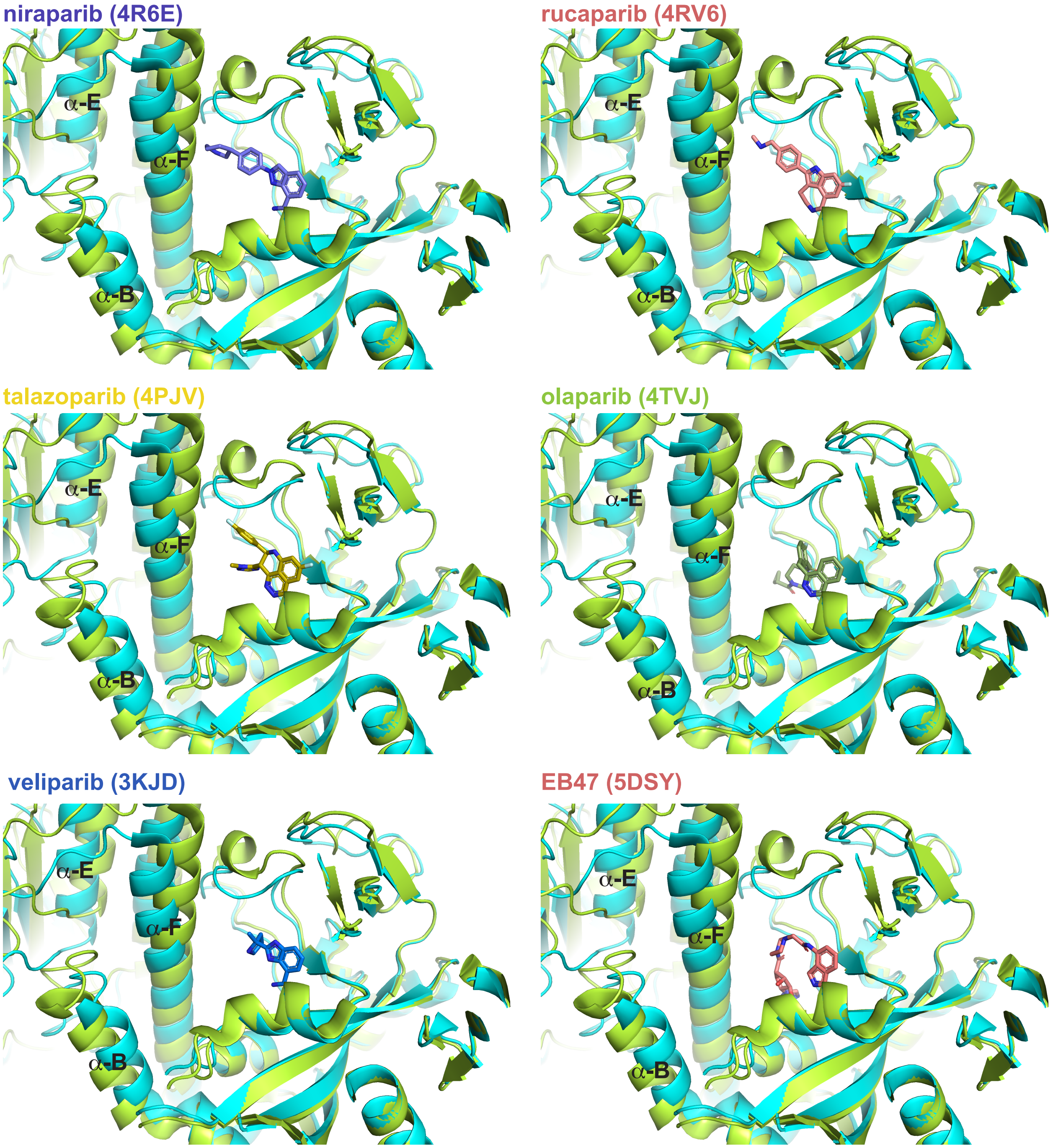


**Fig. S5. Structural alignment of PARP1 and PARP2 bound to DNA showing contacts of various PARPi with helix F.** The crystal structures of PARP1 or PARP 2 CAT domains bound to various PARPi were aligned to the crystal structure of the PARP1/DNA complex (4DQY, teal). The CAT domains from the PARPi complexes are not shown for clarity. The cryo-EM structure of PARP2/HPF1/DNA complex is also overlayed (6X0L, lime) (HPF1 was omitted for clarity).

### SPR binding kinetics

#### PARP2 I318A DMSO

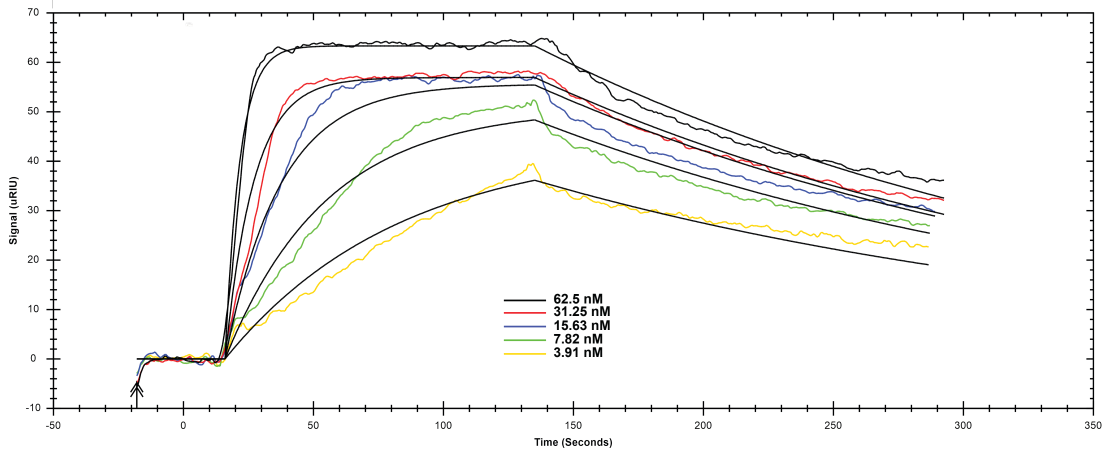


#### PARP2 I318A niraparib

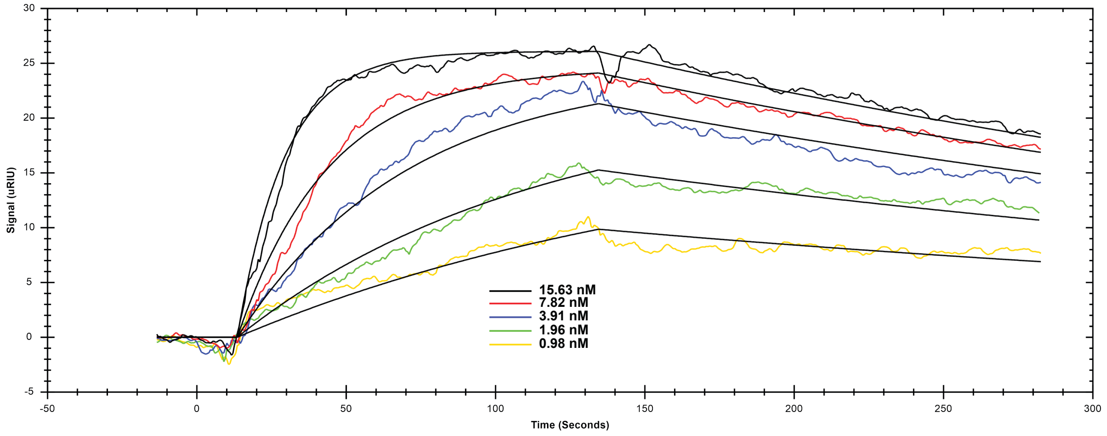


#### PARP2 I318A veliparib

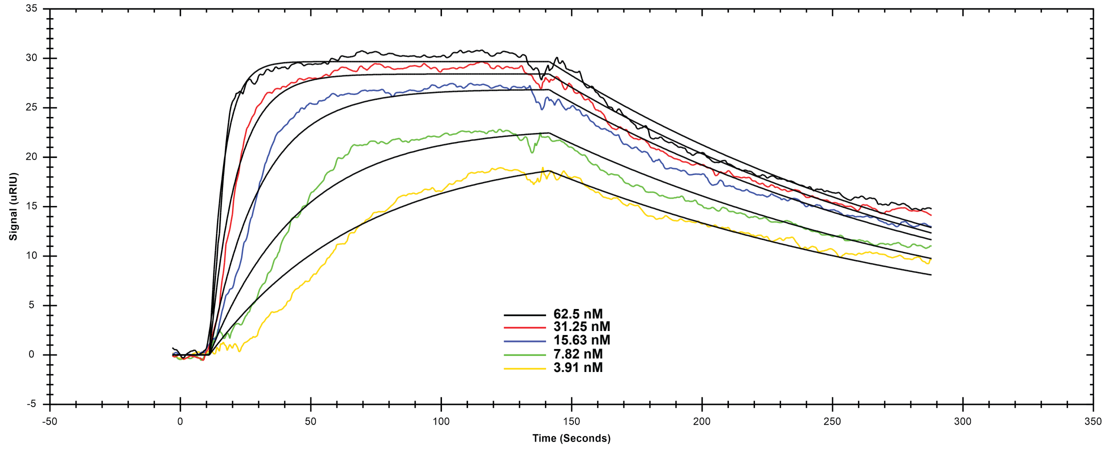

### SPR binding kinetics

#### PARP2 I318V DMSO

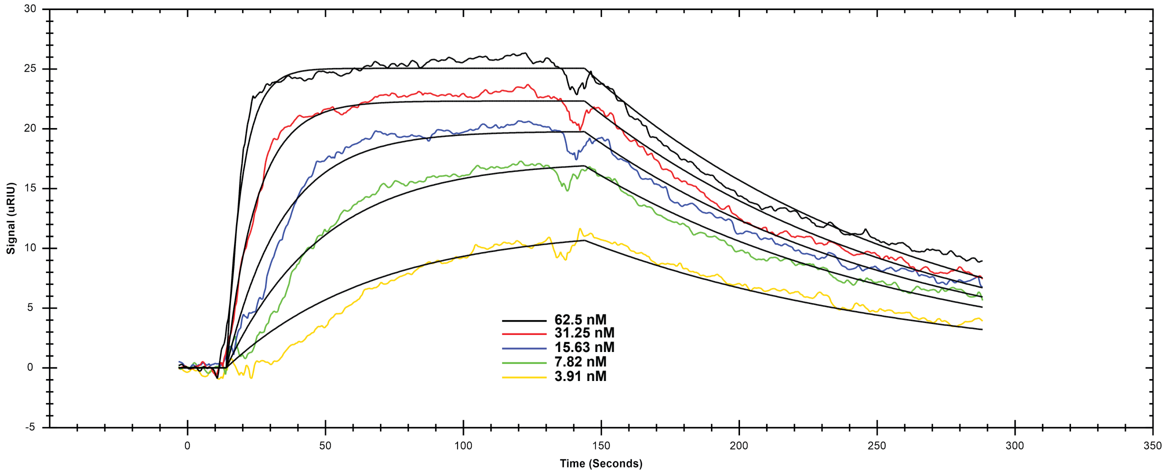


#### PARP2 I318V niraparib

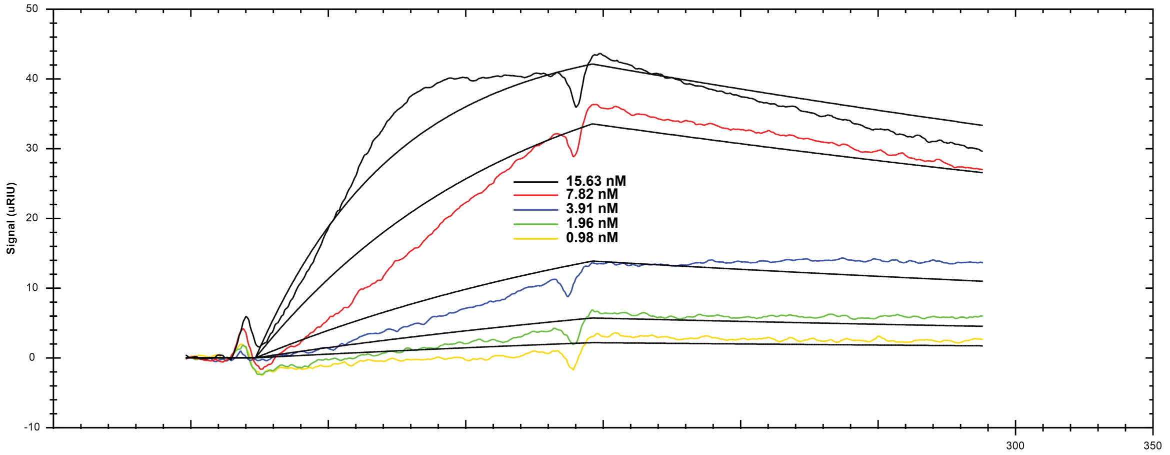


#### PARP2 I318V veliparib

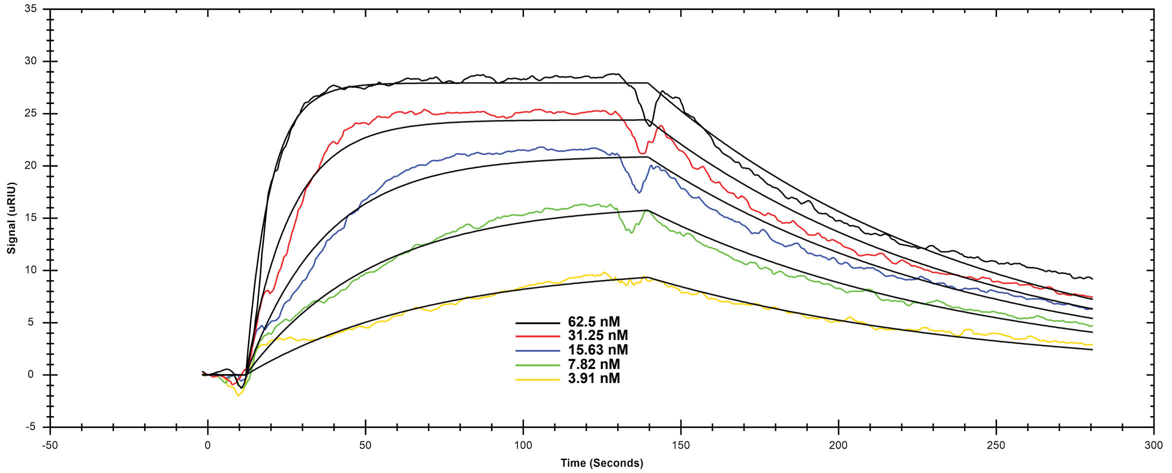

**Fig. S6. SPR binding curve titrations for PARP2 mutants I318A and I318V.** A biotinylated DNA containing a central 5'P nick was captured to a streptavidin coated chip. PARP2 I318A or I318V mutants were flowed at various concentrations over the chip as indicated on the figure in the presence of DMSO or 5 μM inhibitor. A 1:1 binding model was used to fit the data in TraceDrawer (Reichert).

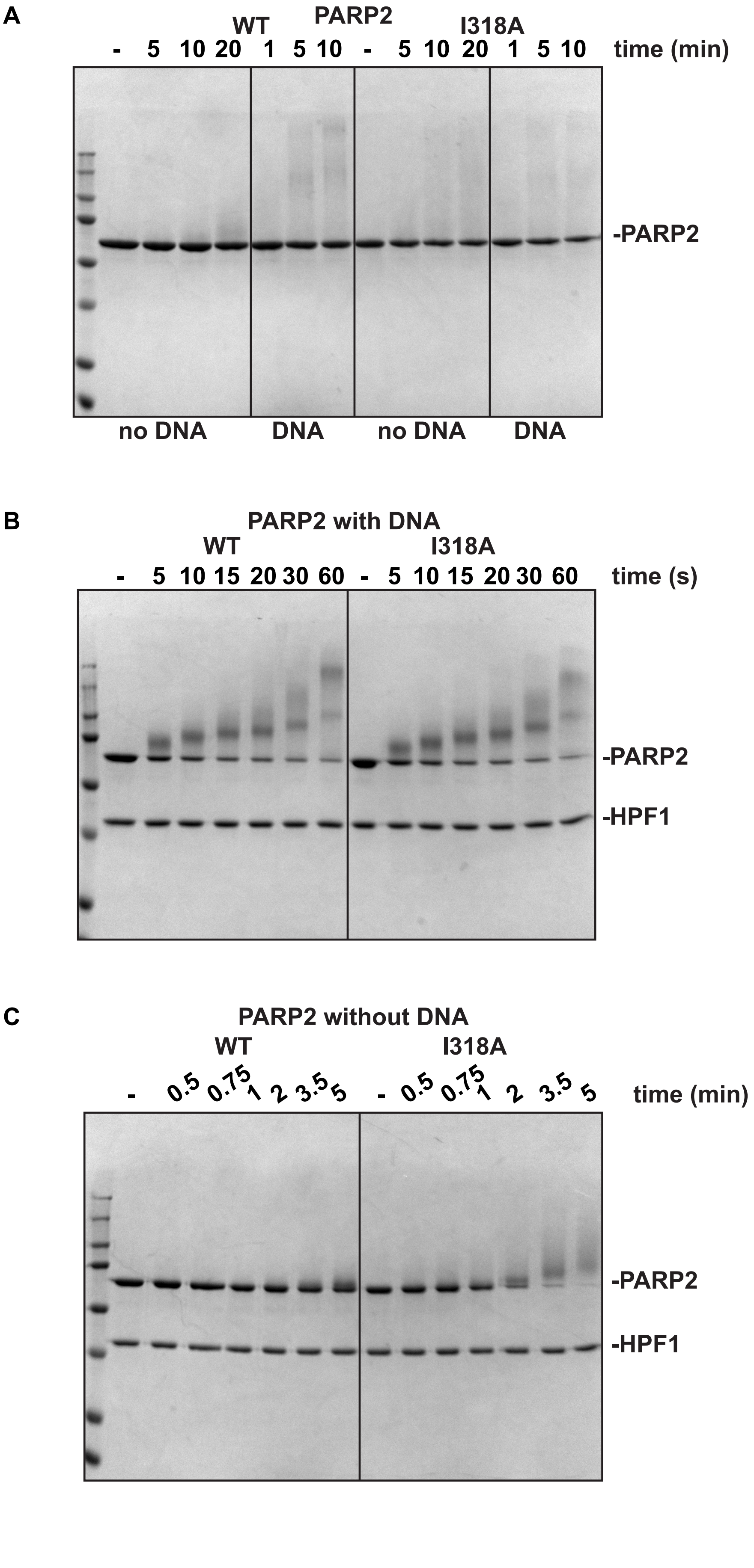
A
PARP2
WT
I318A
- 5 10 20 1 5 10 - 5 10 20 1 5 10
time (min)
-PARP2
no DNA
DNA
no DNA
DNA
B
PARP2 with DNA
WT
I318A
- 5 10 15 20 30 60 - 5 10 15 20 30 60
time (s)
-PARP2
-HPF1
C
PARP2 without DNA
WT
I318A
- 0.5 0.75 1 2 3.5 5 - 0.5 0.75 1 2 3.5 5
time (min)
-PARP2
-HPF1

**Fig. S7. SDS-PAGE activity assay for PARP2 WT and I318A mutant.** (**A**) PARP2 WT or I318A mutant (1 μM) was incubated with or without DNA (1 μM) and $NAD^+$ for various time point as indicated. (**B**) Same as A with DNA and HPF1 (1 μM). (**C**) Same as B without DNA.

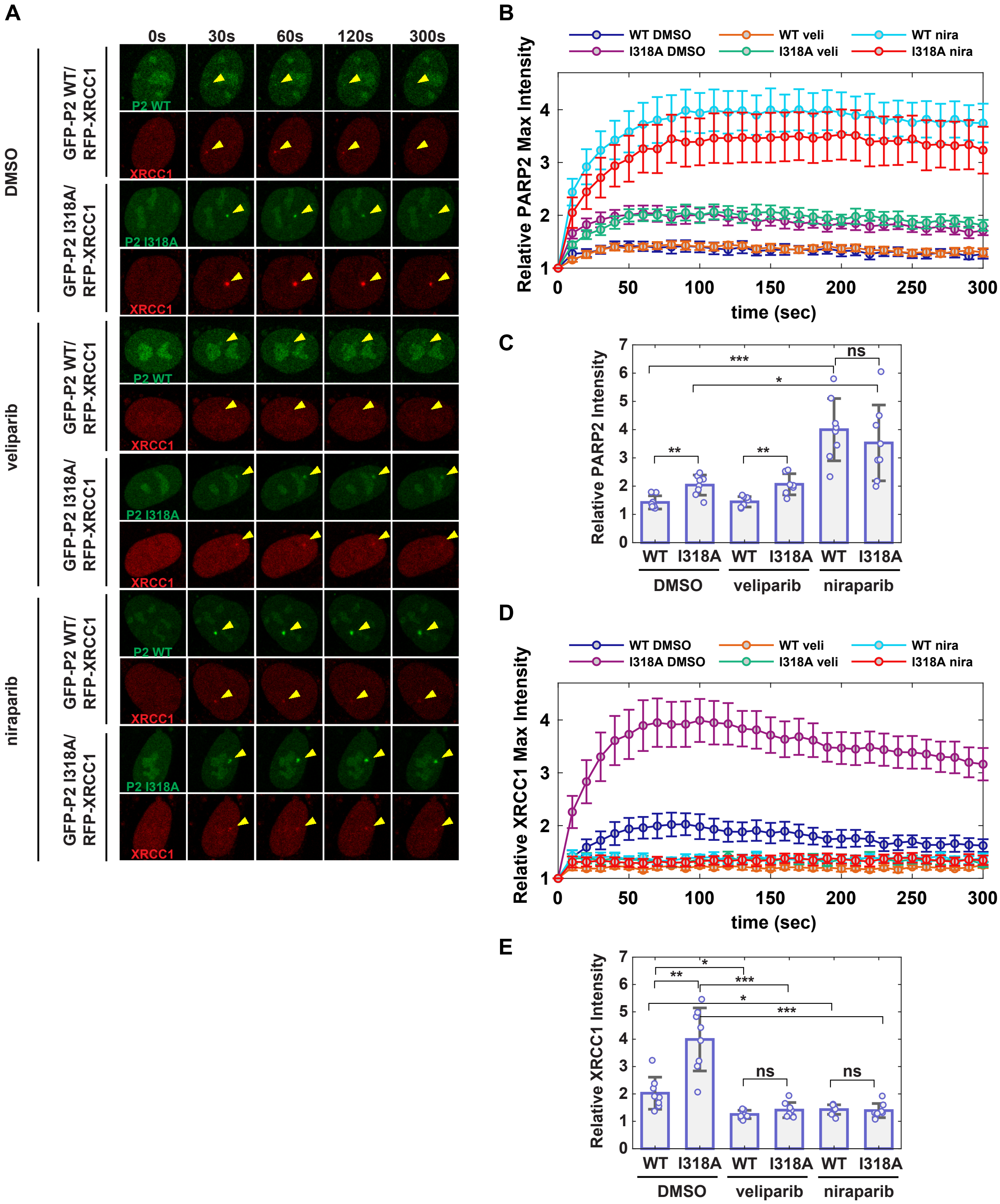


**Fig. S8. PARP2 I318A mutant accumulates more efficiently than PARP2 WT to sites of DNA damage in cells.** (**A**) Representative images of laser-induced GFP-PARP2 WT or I318A mutant and mRFP-XRCC1 foci in PARP1/2 KO RPE-1 cells in the presence of DMSO, veliparib or niraparib. The arrowheads point to the site of micro-irradiation. (**B**) The relative intensity of PARP2 at DNA damage sites (normalized to the intensity before irradiation) in the presence of DMSO, veliparib or niraparib. The points and error bars represent the averages and standard errors, respectively. (**C**) The maximum GFP-PARP2 relative intensity from (**B)**. The bars represent the average of the maximum relative intensity obtained for each cell in one representative experiment out of 3 consistent biological repeats. Each point shown represents the maximum relative intensity obtained for one out of 7-8 cells. The error bars correspond to the standard deviations. Two-sample two-sided *t*-tests were used to compare the relative PARP2 foci intensity between samples as indicated. * asterisks indicate $P < 0.05$, ** asterisks indicate $P < 0.005$, *** asterisks indicate $P < 0.0005$, and ns, indicates not significant. (**D**) and (**E**), the same analysis as (**B)** and (**C)** but for RFP-XRCC1 foci.

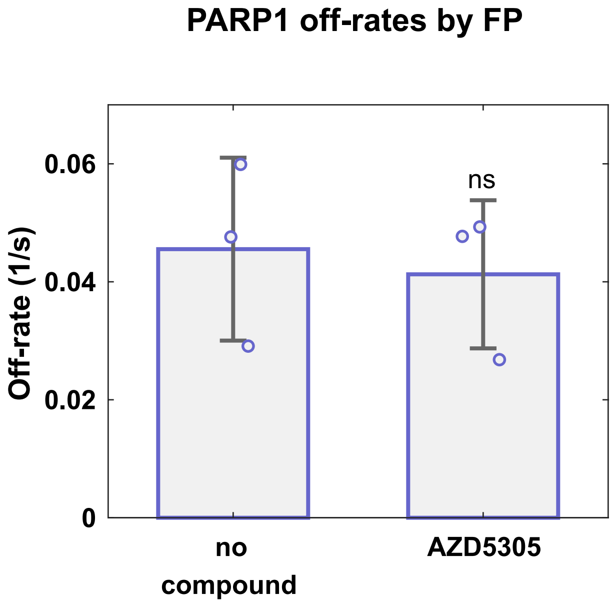


**Fig. S9. Fluorescence polarization DNA competition experiment showing the effect of AZD5305 on PARP1 DNA retention.** PARP1 (40 nM) was incubated with a dumbbell DNA probe containing a central 5'P nick (5 nM) for 30 minutes at room temperature in the presence of DMSO or AZD5305 (100 μM). A competitor unlabelled DNA (2 μM) was added and FP was measured over time. The data was fitted using a single exponential in Matlab to obtain an off-rate. The off-rates shown are an average of 3 independent experiments. The error bar corresponds to the standard deviation. The points represent the off-rate value for each single experiment. Two-sample two-sided *t*-tests were used to compare the off-rate values between samples with AZD5305 and samples with DMSO and ns, indicates not significant.

### FP DNA competition off-rates

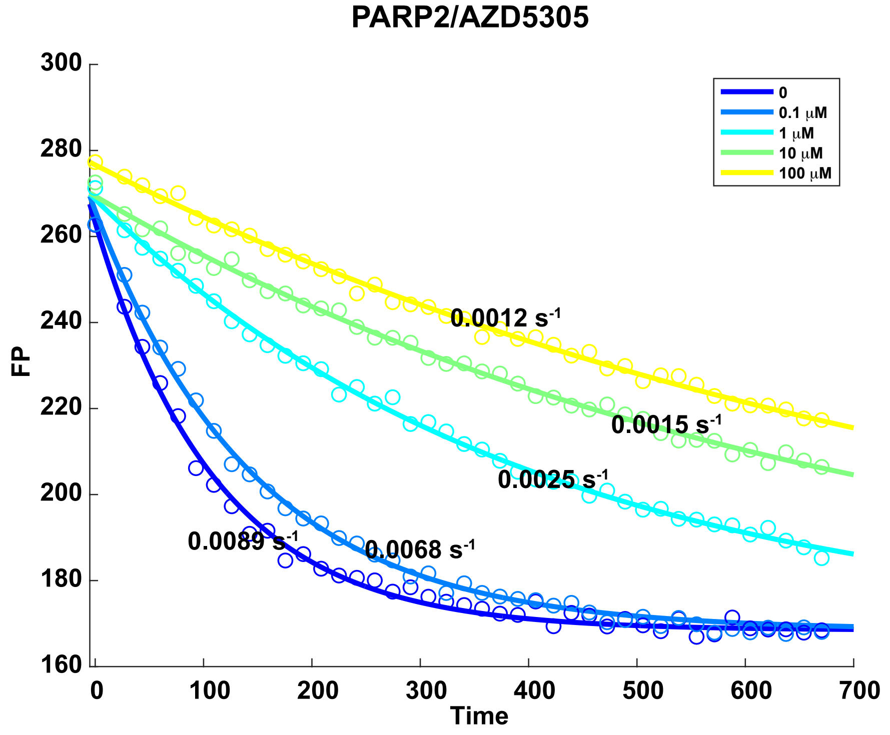


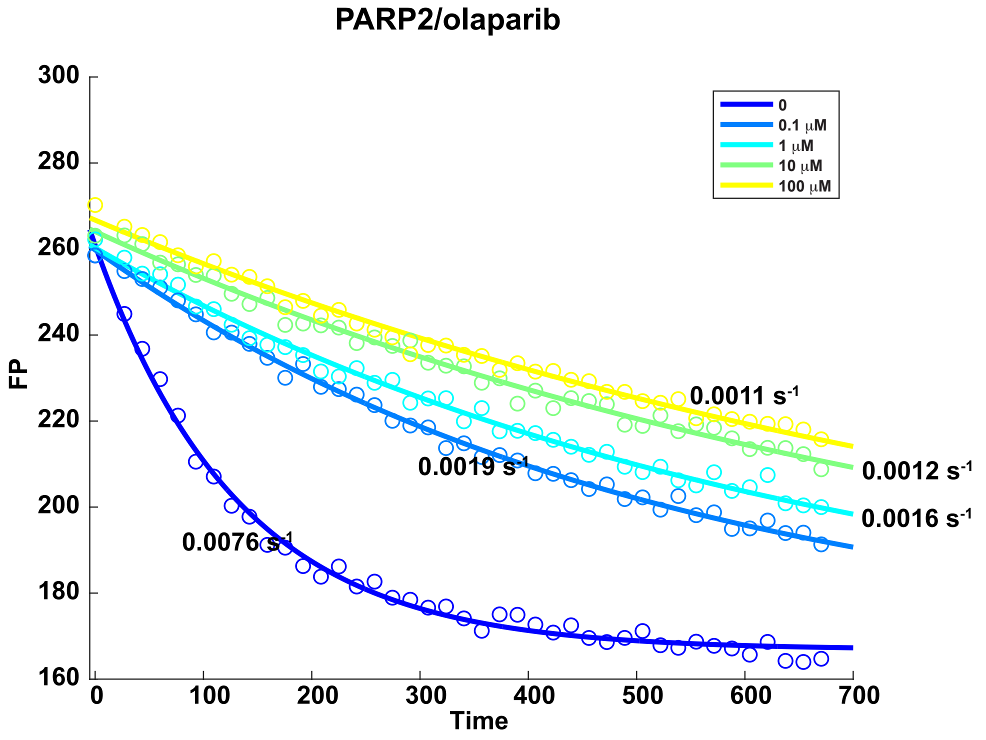

**Fig. S10. Examples of Fluorescence Polarization DNA competition experiments fitting for AZD5305 and Olaparib at various compound concentrations.** PARP2 (40 nM) was incubated with a dumbbell DNA probe containing a central 5'P nick (5 nM) for 30 minutes at room temperature in the presence of AZD5305 or olaparib at various concentrations as indicated. A competitor unlabelled DNA (2 μM) was added and FP was measured over time. The data was fitted to a single exponential in Matlab to obtain an off-rate.
